# Local Delivery of SBRT and IL12 by mRNA Technology Overcomes Immunosuppressive Barriers to Eliminate Pancreatic Cancer

**DOI:** 10.1101/2023.10.30.564833

**Authors:** Angela L. Hughson, Gary Hannon, Noah A. Salama, Tara G. Vrooman, Caroline A. Stockwell, Bradley N. Mills, Jesse Garrett-Larsen, Haoming Qiu, Roula Katerji, Lauren Benoodt, Carl J. Johnston, Joseph D. Murphy, Emma Kruger, Jian Ye, Nicholas W. Gavras, David C. Keeley, Shuyang S. Qin, Maggie L. Lesch, Jason B. Muhitch, Tanzy M.T. Love, Laura M. Calvi, Edith M. Lord, Nadia Luheshi, Jim Elyes, David C. Linehan, Scott A. Gerber

## Abstract

The immunosuppressive milieu in pancreatic cancer (PC) is a significant hurdle to treatments, resulting in survival statistics that have barely changed in 5 decades. Here we present a combination treatment consisting of stereotactic body radiation therapy (SBRT) and IL-12 mRNA lipid nanoparticles delivered directly to pancreatic murine tumors. This treatment was effective against primary and metastatic models, achieving cures in both settings. IL-12 protein concentrations were transient and localized primarily to the tumor. Depleting CD4 and CD8 T cells abrogated treatment efficacy, confirming they were essential to treatment response. Single cell RNA sequencing from SBRT/IL-12 mRNA treated tumors demonstrated not only a complete loss of T cell exhaustion, but also an abundance of highly proliferative and effector T cell subtypes. SBRT elicited T cell receptor clonal expansion, whereas IL-12 licensed these cells with effector function. This is the first report demonstrating the utility of SBRT and IL-12 mRNA in PC.

**Statement of significance:** This study demonstrates the use of a novel combination treatment consisting of radiation and immunotherapy in murine pancreatic tumors. This treatment could effectively treat local and metastatic disease, suggesting it may have the potential to treat a cancer that has not seen a meaningful increase in survival in 5 decades.

## 1. Introduction

Pancreatic ductal adenocarcinoma (PDAC) claims over 450,000 lives globally each year and owns the ominous ranking as the most lethal malignancy of all major cancers [1]. These dismal survival rates can be largely attributed to a signature tumor microenvironment (TME) that fosters immune suppression and imparts treatment resistance [2, 3]. Minimal clinical progress has been made against this malignancy over the past 50 years [4], therefore our laboratory has investigated an alternative treatment strategy that administers two therapies directly to the TME consisting of local radiotherapy and intratumoral immunotherapy. This innovative approach would concentrate the therapeutic payload to the tumor specifically, which we hypothesized could overcome the immunosuppressive properties of the PDAC TME.

Stereotactic body radiotherapy (SBRT) has largely replaced conventional radiotherapy during the past decade, emerging as a viable treatment adjunct for localized PDAC [5, 6]. SBRT enables precise delivery of high-dose radiation (20–35 Gy total) over a short period of time (5–6 fractions in 1–2 weeks), which is convenient for patients and causes minimal radiation damage to surrounding tissues [5]. Our recent clinical study demonstrated that SBRT administered to human PDAC *initiated* an antitumor response, but this response was restrained by both existing and SBRT-induced suppressive mechanisms [7]. Since it is well established that the immune system mediates many of the antitumor effects of radiotherapy [8], we investigated whether a combination therapy targeting the immune system would enhance the efficacy of radiotherapy in PDAC.

Interleukin-12 (IL-12) is a proinflammatory, pleiotropic cytokine with potent activity on both the adaptive and innate immune systems [9]. IL-12 was initially investigated as an anticancer agent due to its ability to enhance the cytotoxic function of effector T cells and repolarize suppressive myeloid cells. Unfortunately, systemic administration resulted in toxicity and produced only modest outcomes [10, 11]. Here, we utilized a novel method for intratumoral delivery of murine IL-12 using mRNA technology. This strategy consists of a unique IL-12p70 mRNA sequence where the p35 and p40 heterodimeric subunits were linked by a polypeptide linker resulting in a bioactive IL-12p70 fusion protein [12]. This construct is encapsulated in a lipid nanoparticle to facilitate cellular uptake and release of the mRNA from the endosome, all while protecting the nucleic acid from destruction. This represents the first report of its kind combining this IL-12mRNA technology with SBRT in PDAC.

SBRT/IL-12 therapy resulted in remarkable tumor control and survival in preclinical orthotopic and spontaneous PDAC models of primary and metastatic disease. Intratumoral concentrations of IL-12 remained local and were augmented by SBRT. Combined therapy led to a complete loss of exhausted T cells and the presence of an activated, effector T cell phenotype that also displayed robust intratumoral proliferation. SBRT caused clonal expansion of the TCR repertoire whereas IL-12 induced T cell IFNγ production, which was required for treatment efficacy.

## 2. Results

### 2.1 Local delivery of SBRT and IL-12mRNA therapy eliminates primary disease and promotes durable, systemic antitumor immunity

We developed a model where both SBRT and immunotherapy were delivered locally to the primary pancreatic tumor (**Figure 1A**). Mice were injected orthotopically with the luciferase-expressing KP2 cell line and established tumors were treated with a clinically relevant dose and schedule of SBRT consisting of 6Gy/day for 4 consecutive days. One day following the final fraction of SBRT, an intratumoral injection of IL-12mRNA or scrambled control mRNA (scRNA; non-translating) was administered and tumor burden was monitored by luciferase-detecting IVIS along with survival. This regimen resulted in four treatment groups: 1) No SBRT + scRNA (hereafter referred to as “Untreated”), 2) SBRT + scRNA (“SBRT”), 3) No SBRT + IL-12mRNA (“IL-12mRNA”), and 4) SBRT + IL-12mRNA (“SBRT/IL-12mRNA”). SBRT/IL-12mRNA was superior to each monotherapy and resulted in a dramatic reduction of primary tumor burden (**Figure 1B**) including long-term survival (**Figure 1C**) past 300 days. We analyzed the rejection kinetics on a per tumor basis and demonstrated a varied decrease in tumor size as early as 24-48 hours post-IL-12mRNA injection, with all mice reverting to baseline BLI between 3– and 13-days following completion of treatment (**Supplemental Figure 1; see grey shaded region**). Cured mice rested for 6 weeks following SBRT/IL-12mRNA treatment were able to reject a hepatic challenge of PDAC demonstrating long-term systemic immunity (**Figure 1D**). The aggressive KP2 cell line used in the experiments described above has driver mutations in *KRAS* and *TP53*; a phenotype that accounts for approximately 70% of human PDAC patients [19]. We tested whether SBRT/IL-12mRNA was effective in other variants of PDAC, including mutated *KRAS* only (KCKO cell line) and the highly aggressive and incurable spontaneous KPC (LSL-Kras^G12D^; Tp53^L/L^; P48-Cre) mouse model that develops tumors throughout the entire pancreas [13]. Similar to the KP2 model, SBRT/IL-12mRNA was highly efficacious against KCKO tumors (**Supplemental Figure 2A, B, C**) and provided a significant survival advantage in homozygous KPC mice (**Supplemental Figure 2D**).

**Figure 1:**
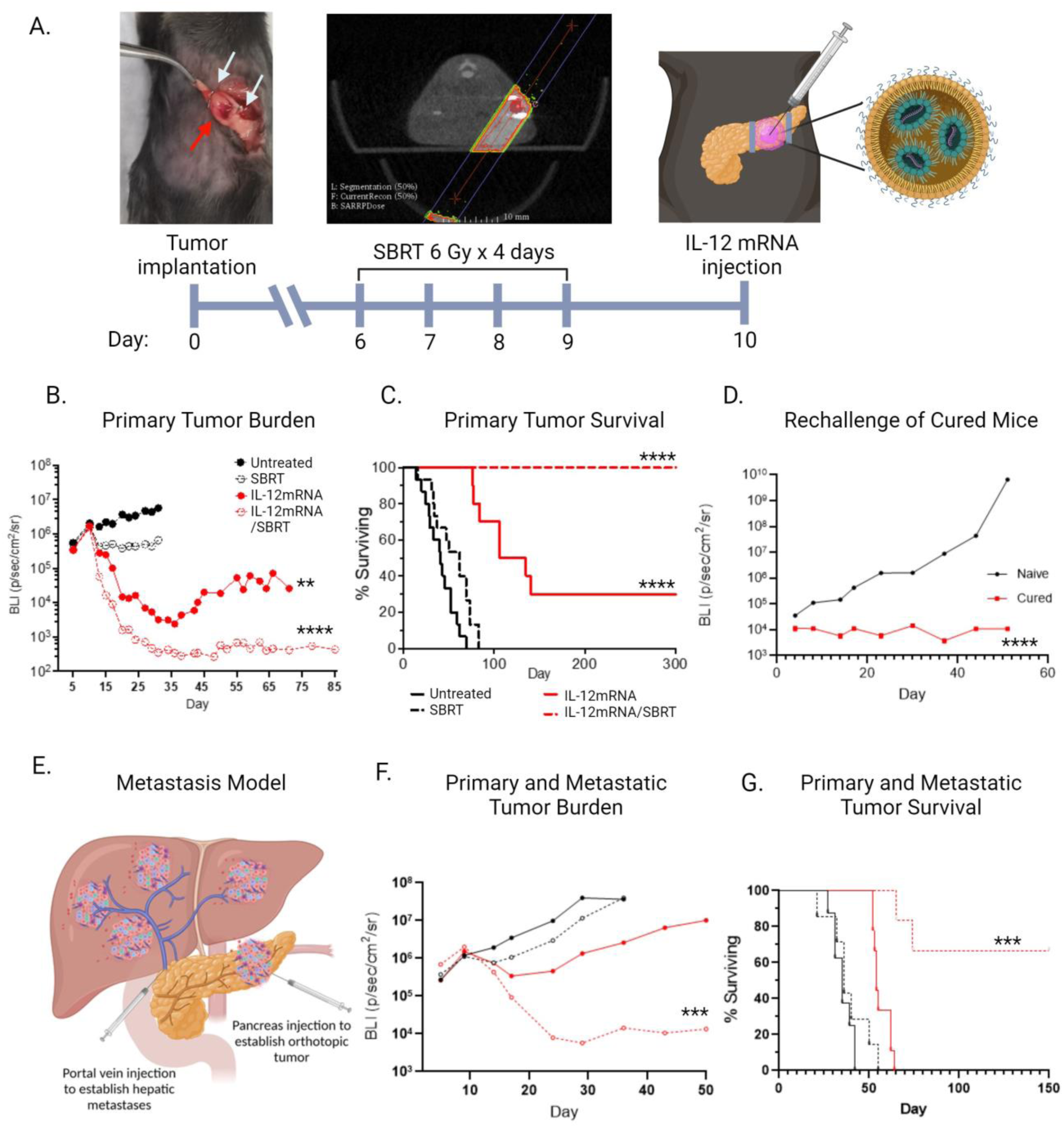
Combination treatment of SBRT + IL-12mRNA eliminates pancreatic tumors and promotes survival. (**A**) Illustration of PDAC mouse model where 2.5×10^4^ KP2-luciferase-expressing tumor cells were injected orthotopically (red arrow) and flanked with titanium fiducial clips (white arrows) for SBRT targeting. The tumors were treated with or without SBRT (6G/day over 4 consecutive days) followed by an intratumoral injection of IL-12mRNA or control scRNA 1 day after the final fraction of radiation. (**B**) Tumor burden for primary disease was assessed by weekly BLI measurements for each treatment. (**C**) Survival was tracked out to 300 days post tumor injection. (**D**) BLI growth curve for systemic rechallenge of cured mice versus aged-matched naïve mice (**E**) Illustration of the metastatic model developed to test the systemic effects SBRT/IL-12mRNA treatment: pancreas and hepatic portal vein were injected concurrently to establish primary (pancreas) and metastatic (liver) tumors. (**F**) BLI growth curve for each treatment for the metastatic PDAC model. (**G**) Survival curve for the metastatic model was tracked out to 150 days. n=15/group from 3 independent experiments in B and C. n=5 for cured mice and n=9 aged-matched naïve mice in D. n=6-8 in F and G. For BLI growth curves, mean values for each time point are presented for each treatment group. **p<0.01; p<0.0001.

To test the systemic effects of our therapy, we generated a clinically relevant metastatic model where localized therapy was administered to the primary tumor alone to determine whether distal effects could be achieved. Mice were injected with primary orthotopic pancreatic tumors as illustrated in **Figure 1A** and concurrently given a portal vein injection to recapitulate distal hepatic metastases; the most common site for PDAC dissemination (**Figure 1E**). Even though only the primary tumor received SBRT/IL-12mRNA treatment, significant distal effects to liver metastasis were observed that corresponded to long-term survival in 67% of mice (**Figure 1F & G**). Similar systemic antitumor immunity was also observed using the KCKO metastatic model (**Supplemental Figure 2E & F**). Collectively, these data demonstrate that localized delivery of SBRT and IL-12mRNA eliminates primary PDAC tumors and promotes systemic immunity capable of controlling distal metastatic disease.

### 2.2 IL-12mRNA administered to the TME results in local concentrations of IL-12 protein

We quantified the concentration of IL-12 protein following intratumoral administration of IL-12mRNA and whether this cytokine was confined locally. As illustrated in **Supplemental Figure 3A**, SBRT/IL-12mRNA results in the highest production of intratumoral IL-12 at 24 hours, which returns near baseline at 96 hours. Similar concentrations and kinetics were detected in the pancreatic draining lymph node (dLN) (**Supplemental Figure 3B),** however near 20-fold lesser concentrations were measured in distal tissues such as the liver and non-draining lymph node (ndLN) at 24 hours, and no IL-12 protein above controls was detected at 96 hours (**Supplemental Figure 3C & D**). The production of IL-12 protein suggests that IL-12mRNA was being taken up and translated to protein by cells in the TME. We performed *in vitro* assays to identify which cells were responsible for this production. Cell subsets commonly found in the PDAC TME (tumor cells, fibroblasts, lymphocytes, and macrophages) were cultured with either IL-12mRNA or scRNA sequence and qPCR was used to determine mRNA uptake by detecting a unique RNA sequence located in the IL-12mRNA. All cells analyzed were able to take up abundant IL-12mRNA when compared to scRNA controls (**Supplemental Figure 3E**), however only fibroblasts and especially tumor cells and macrophages produced IL-12 protein following uptake (**Supplemental Figure 3F**). These data demonstrate that intratumoral administration of IL-12mRNA results in elevated local concentrations of IL-12 protein in the TME likely produced by tumor and host stromal/myeloid cells.

### 2.3 Analysis of the TME following SBRT/IL-12mRNA

The PDAC TME is highly immunosuppressive and represents a major barrier to treatment efficacy. We assessed whether SBRT/IL-12mRNA therapy modulated the TME. Local delivery of SBRT/IL-12mRNA did not alter the percentage of intratumoral CD45+ immune cells as assessed by flow cytometry (Gating scheme defined in **Supplemental Figure 4**) at 24– and 72-hours post IL-12mRNA treatment (**Supplemental Figure 5A**). A flow cytometric immune subset analysis revealed a variety of intratumoral immune cells with macrophages occupying the largest density (**Supplemental Figure 5B**). We observed a reduction of radiosensitive CD4 and B lymphocytes (**Supplemental Figure 5C**). At 72 hours, most of these changes returned to control levels with only slight increases in CD4+ T cells and monocytes persisting in the IL-12mRNA and SBRT/IL-12mRNA treated groups. Histological assessment of the TME by a blinded board-certified pathologist revealed a poorly differentiated carcinoma surrounded by desmoplastic stroma and mild inflammatory infiltrate in all groups (**Supplemental Figure 5D**). Of note were minor increases of fibrosis in the SBRT only group, no definitive treatment response in the IL-12mRNA only group, but appreciable treatment response in the SBRT/IL-12mRNA group defined as fibrosis, foci of necrosis, hyalinization, and mixed inflammatory infiltrate. Collectively, although the tumor immune cell infiltrate was largely similar between all groups, only SBRT/IL-12mRNA demonstrated notable pathological changes in the TME.

### 2.4 T cells are essential to mediate SBRT/IL-12mRNA efficacy

We investigated which cell types were responsible for mediating the marked tumor control observed following SBRT/IL-12mRNA treatment using antibody depletion of the effector cell populations (CD4 and CD8 T cells; NK cells). NK cell depletion had no effect on tumor burden or survival (**Supplemental Figure 6A**). However, 50% of SBRT/IL-12mRNA efficacy was lost when either CD4 or CD8 T cells were depleted individually (**Supplemental Figure 6B & C** respectively), including a complete loss of effectiveness when CD4 and CD8 T cells were depleted concurrently (**Supplementary** Figure 6D). These data identify T cells as the effector population responsible for PDAC elimination following SBRT/IL-12mRNA therapy.

### 2.5 Transcriptomic analysis of the TME following SBRT/IL-12mRNA therapy

Since no appreciable differences were seen in the quantity of intratumoral immune cells following therapy (**Supplemental Figure 5A & B**), we hypothesized SBRT/IL-12mRNA may enhance the quality of these cells. Single cell RNA-sequencing (scRNA-seq) was utilized to assess transcriptomic differences induced by SBRT/IL-12mRNA. Following quality control and filtering steps, over 11,000 intratumoral cells from each treatment group were sequenced as detailed in the materials and methods. Unsupervised clustering of all cells identified 29 distinct clusters, which were further annotated into distinct cell subsets based on canonical markers and assembled into four larger groups namely tumor/stromal cells, myeloid cells, B cells, and T/NK cells (**Figure 2A**). Individual Uniform Manifold Approximation and Projection (UMAPs) plots were generated for each treatment condition (**Figure 2B**) and the relative position of 4 major cellular groups was assessed between each treatment. The myeloid and T/NK cell groups demonstrated appreciable positional movement through the UMAP as conditions progressed from Untreated to SBRT/IL-12mRNA suggesting transcriptomic differences attributed to therapy. Therefore, we focused our analysis on these two intratumoral cell subsets.

**Figure 2:**
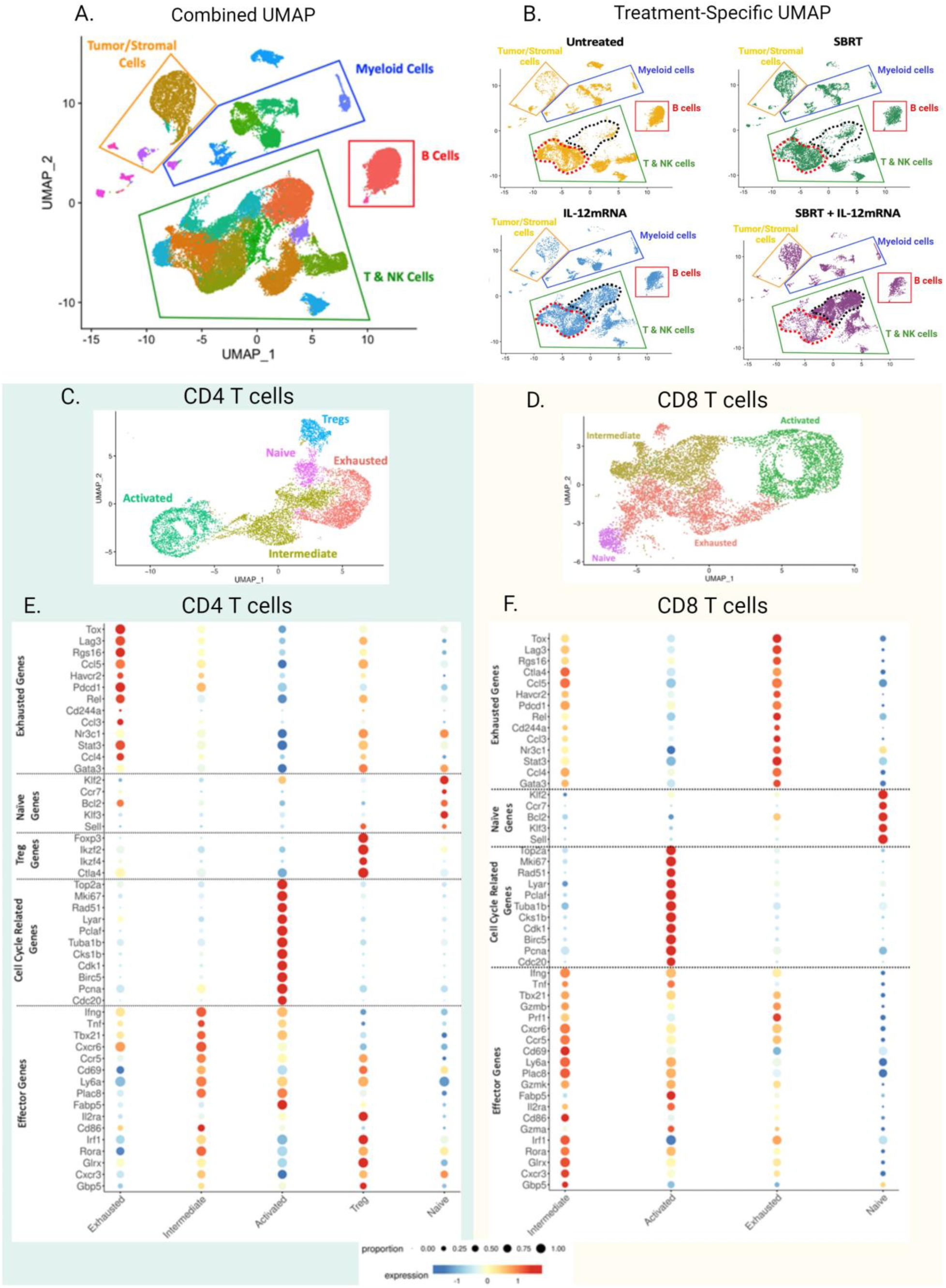
T cells undergo distinct transcriptomic changes following SBRT/IL-12mRNA treatment. (**A**) Unsupervised clustering of intratumoral cells following scRNA-seq could identify 4 distinct cell subsets: Tumor/Stromal cells, Myeloid cells, B cells and T/NK cells. (**B**) Separating this clustering to treatment-specific UMAPs could identify significant transcriptomic changes to T cells, in particular, following SBRT/IL-12mRNA treatment. (**C** & **D**) Subclustering on CD4 and CD8 T cells specifically identified populations of naïve, exhausted, intermediate and activated for each. (**E & F**) Dot plots highlight expression levels of hallmark genes for a given proportion of cells associated with each subcluster. See materials and methods for further details.

The immunostimulatory/suppressive phenotype of intratumoral myeloid cells was assessed by single cell RNA sequencing (scRNA-seq). Myeloid populations were subclustered, and based on canonical markers, resulted in predominate populations of dendritic cells (DCs), macrophages and monocytes, and neutrophils (**Supplemental Figure 7**). SBRT/IL12mRNA resulted in heightened expression of many immunostimulatory genes when either all myeloid cells or each individual subset were examined against Untreated and SBRT only groups (**Supplemental Figure 8**). This phenotype was largely attributed to IL-12 as the treatment groups consisting of IL-12mRNA only and SBRT/IL-12mRNA clustered together when assessed in an unbiased manner (see clustering brackets atop each map in **Supplemental Figure 8**). These data illustrate the repolarization of intratumoral myeloid cells populations following SBRT/IL-12mRNA therapy.

Cells within the T/NK cell green quadrilateral demonstrated the most visible UMAP changes (**Figure 2B**; note T cells reside in red outline in Untreated, but progress into black outline in SBRT/IL-12mRNA). These data emphasize the greatest impact of SBRT/IL-12mRNA on the T cell subset and complement our depletion data, which also identified T cells as the chief cells responsible for treatment efficacy. Given these results, we directed our analysis on subclustered T cells based on expression of CD4 and CD8 along with coexpression of CD3ε and TCRα/β gene signatures. Utilizing canonical markers, differentially expressed genes (DEGs) (**Supplemental Figure 9A & B**), and curated gene signatures, we identified 5 distinct CD4 T cell subsets (Naïve, T-regulatory, Exhausted, Intermediate, Activated/Cycling) (**Figure 2C**) and 4 CD8 T cell subsets (Naïve, Exhausted, Intermediate, Activated/Cycling) (**Figure 2D**). Naïve and T-regulatory (CD4) subsets encompassed a relatively small percentage of total T cells. The much larger population of Exhausted T cells exhibited abundant expression of the hallmark *TOX, LAG3, HAVCR2, PDCD1, and RGS16* genes (**Figure 2E & F**). Interestingly, the Activated/Cycling population predominately expressed genes associated with cell division (*TOP2A, MKI67, RAD51, PCNA, etc*.), with only moderate expression of classical effector genes such as *IFNG GZMB, PRF1, TBX21, etc.* This is in contrast to the Intermediate subset, which had a far lesser proliferative signature, but heightened expression of effector genes (**Figure 2E & F**). Collectively, T cells mediate SBRT/IL-12mRNA efficacy for which we identified distinct larger subpopulations such as Exhausted, Intermediate, and Activated/Cycling that underwent profound transcriptomic changes following treatment.

### 2.6 SBRT/IL-12mRNA remodels T cell subsets toward an effector/activated phenotype

Individual UMAPs were generated for each treatment condition to parse out how each therapy was modulating T cell subsets. The vast majority of CD4 (95%) and CD8 (88%) T cells in Untreated tumors exhibited transcriptomic signatures of Exhaustion (red population), Regulatory (CD4; purple population), or Naïve (blue population) (**Figure 3A and B** respectively) T cells. These results are in accordance with clinical PDAC data where most intratumoral T cells are dysfunctional and have an impaired ability to respond adequately against the tumor [20]. Although each monotherapy reduced the Naïve and Exhausted populations and increased the Intermediate subsets, SBRT/IL-12mRNA resulted in the noted disappearance of Exhausted/Naïve/Regulatory T cells and a significant expansion of the Activated/Cycling and Intermediate CD4 and CD8 T cell populations (**Figure 3A and B**; note loss of red cells and increase of green and gold populations). Additionally, while IL-12mRNA alone led to an expansion of activated/cycling CD4 and CD8 T cells, these cells still maintained significantly higher expression of immunosuppressive transcripts (*TOX, HAVCR2, RGS16, LAG3, PDCD1, CTLA4*) when directly compared to the activated/cycling cells in the SBRT/IL-12mRNA treated group (**Supplemental Figure 10A**).

**Figure 3.**
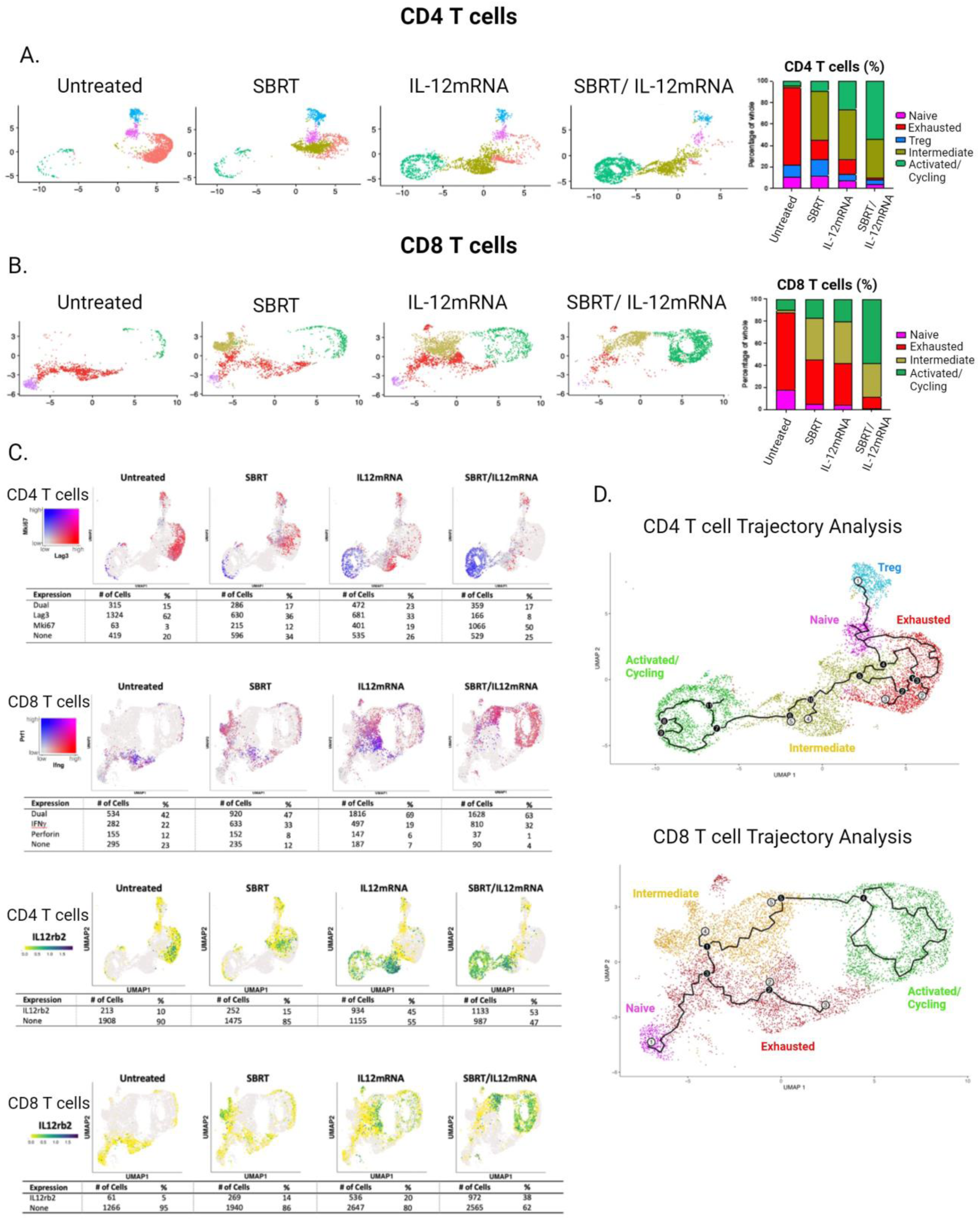
SBRT/IL-12mRNA treatment remodels the CD4 and CD8 T cell response in tumors. (**A** & **B**) Subclustered UMAPs and relative proportions of CD4 and CD8 T cell subpopulations following each treatment. (**C; top and middle**) Co-expression heatmaps for Mki67/Lag3 for CD4 T cells and Prf1/Ifng for CD8 T cells for each treatment. (**C; bottom**) IL12rb2 expression on CD4 T cells and CD8 T cells for each treatment. (**D**) Trajectory analysis for CD4 and CD8 T cells with Naïve T cells utilized as the root population in each case. Black circles indicated branch nodes and white circles signify potential terminal outcomes along the trajectory. See materials and methods for further details.

Further analysis of the Exhausted CD4 T cell subpopulation by coexpression feature plots revealed the majority of *LAG3*-expressing cells (81%) did not express the proliferative marker *MKI67* in Untreated tumors (**Figure 3C; top**). This is in contrast to SBRT/IL-12mRNA tumors where not only was there a marked reduction of *LAG3*-expressing T cells, but the proliferative signature shifted towards the Activated/Cycling population. Similar data was observed in CD8 T cells (**Supplemental Figure 10B**).

We next examined the effector markers IFNγ and perforin (*PRF1*) to investigate whether SBRT/IL-12mRNA modulates the cytotoxicity profile of CD8 T cells. Treatment results in a complete conversion (96% of CD8) towards cells that either express IFNγ (32%) or both IFNγ and perforin (63%) suggestive of cytotoxic effector cells (**Figure 3C; middle**).

IL-12 must bind its cognate receptor, which is comprised of the IL-12rb1 and IL-12rb2 subunits, to elicit direct cellular effects. We assessed the expression of these subunits in the CD4 and CD8 T cell populations with and without therapy. **Figure 3C, bottom two** panels illustrate that the addition of IL-12, either alone or especially in combination with SBRT, induces expression of IL-12rb2 (yellow signifies low to no expression whereas green indicates upregulation) and IL-12rb1 (**Supplemental Figure 10C & D**). These data suggest that increasing the intratumoral concentration of IL-12 may elicit a positive feedforward effect allowing intratumoral T cells to be more responsive to this cytokine.

Intratumoral T cells can transition between different functional states (*e.g.* exhausted, activated, etc.), based on the type of therapy. We performed unsupervised Monocle 3 pseudotime inference to assess how SBRT/IL-12mRNA modulates T cell trajectories. Black circles indicated branch nodes, where cells can travel to one of several outcomes, whereas white circles denote a potential terminal outcome of the cell trajectory. CD4 T cell pseudotime analysis using Naïve T cells as the root population identified several trajectories depending on condition: Tregs (outcome 1), Exhausted T cells (outcome 2, 3), and the Intermediate T cell population (outcome 4, 5) (**Figure 3D, top**). Importantly, the CD4 Exhausted and Intermediate T cells were neighboring clusters connected by branchpoints along the pseudotime trajectories suggesting a possible transition between these two states. The Activated/Cycling CD4+ T cell population did not have terminal outcome nodes and was only connected to the Intermediate cluster suggesting that Activated/Cycling cells could give rise to the Intermediate population or that only the Intermediate population could enter the Activated/Cycling state. Similar results were observed in the CD8 T cell trajectory analysis where terminal outcome nodes only existed in the Exhausted and Intermediate subsets, but not the Activated/Cycling population (**Figure 3D, bottom**). Similar to CD4 T cells, a link also existed between the CD8 Intermediate population and the Activated/Cycling cells, but not between the Exhausted and Activated/Cycling populations.

Collectively, these data demonstrate that the Exhausted T cell subset observed almost exclusively in Untreated tumors is largely absent following SBRT/IL-12mRNA therapy and instead replaced with T cells exhibiting an activated/effector phenotype.

### 2.7 SBRT/IL-12mRNA induces intratumoral T cell proliferation

Cell cycle/proliferation was one of the strongest transcriptomic signatures observed in T cells following SBRT/IL-12mRNA. To further investigate this finding, we performed a Cell Cycle Score, which assesses expression of genes associated with different stages of cell cycle (G1; S-phase; G2/M), on both intratumoral CD4 (**Figure 4A**) and CD8 (**Figure 4B**) T cells. For both CD4 and CD8 T cells, the majority of Naïve, Exhausted, and Intermediate cells exhibited a “G1” signature suggesting a phenotype of minimal proliferation. In contrast, the Activated/Cycling T cells formed a circular cluster with approximately one-half the cells in S-phase and the other half in G2/M. This strong proliferative signature shaped the UMAP essentially into dividing (cells in S-phase and G2/M) and non-dividing (G1) T cells.

**Figure 4.**
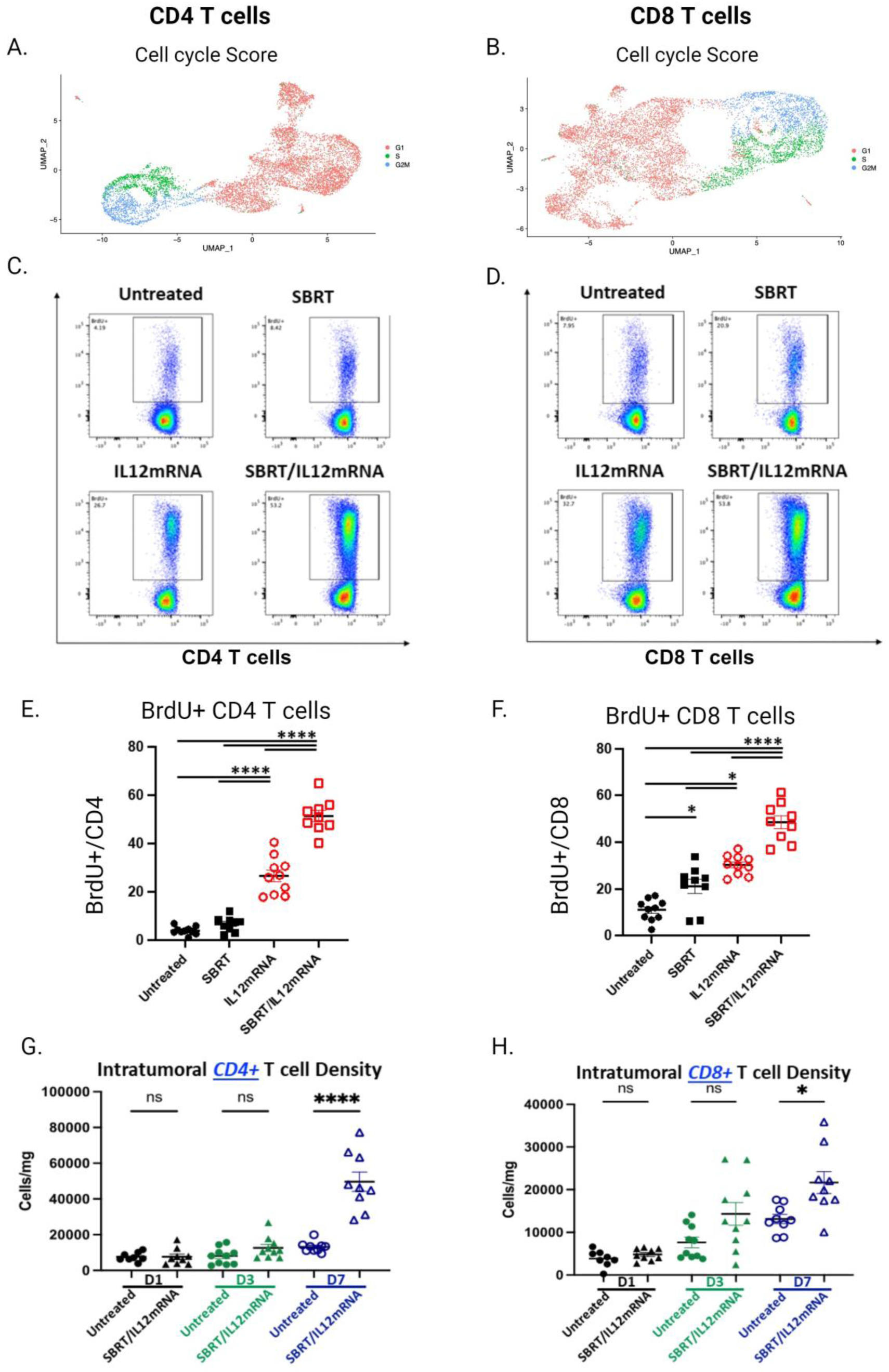
SBRT/IL-12mRNA induces proliferation in tumor CD4 and CD8 T cells. (**A** & **B**) Cell cycle score for CD4 and CD8 T cells using genes associated with G1, S, and G2/M phases of cell cycle. (**C** & **D**) Representative flow graphs for BrdU staining in CD4 and CD8 T cells at 3 days post IL-12mRNA/scRNA injection. (**E** & **F**) Quantification of BrdU+ CD4 and CD8 T cells. (**G** & **H**) CD4 and CD8 T cell tumor levels at day 1, day 3 and day 7 post IL-12mRNA/scRNA injection. n=10/group from 2 independent experiments in C – H. One-way ANOVA followed by Tukey’s test. Significance against untreated and between treatments are reported. *P<0.05; ****P<0.0001.

To validate these transcriptomic data, mice bearing KP2 tumors and treated as described in **Figure 1A** were administered BrdU 4 hours before sacrifice (3 days post IL-12mRNA/scRNA treatment). BrdU incorporation signifying proliferating intratumoral T cells was assayed by flow cytometry. As illustrated in **Figure 4C & D** and quantified in **Figure 4E & F**, CD4 and CD8 T cells from untreated tumors had minimal BrdU incorporation, whereas T cells from tumors treated with SBRT/IL-12mRNA exhibited significant BrdU incorporation suggestive of robust proliferation. Approximately one-half of all intratumoral T cells were dividing, which strongly supports the transcriptomic results demonstrating increased expression of cell cycle/proliferation genes following SBRT/IL-12mRNA therapy.

Robust proliferation induced by SBRT/IL-12mRNA treatment resulted in an increased density of both intratumoral CD4 and CD8 T cells as determined by flow cytometry (**Figures 4G and H** respectively). The elevated density of T cells was not immediate, but rather delayed as significant increases were not observed until 7 days after IL-12mRNA administration. Collectively, these data demonstrate that the combination of SBRT with IL-12mRNA results in vigorous cell division of intratumoral T cells, which in turn augments T cell density.

### 2.8 SBRT/IL-12mRNA results in a clonal TCR repertoire, which is largely a function of radiotherapy

Radiotherapy, through the release of antigen or provision of neoantigens, has been shown to remodel the TCR repertoire [21, 22]. We investigated whether SBRT alone, or in combination with IL-12mRNA, modulates the repertoire by performing TCR sequencing on intratumoral T cells isolated from tumors 3 days after IL-12mRNA or scRNA administration (**Figure 5A**). We assessed the number of unique clonotypes of each treatment group using the well-established Chao1 Diversity/Clonality Score Index with downsampling to normalize input. Both CD4 and CD8 T cells from the SBRT and SBRT/IL-12mRNA groups exhibited a more clonal TCR repertoire when compared to Untreated and IL-12mRNA (**Figure 5B**). This analysis considers the entire repertoire, therefore we focused on the top 1%, which likely represents the more dominant antitumor TCR clonotypes. Bubbleplots of these top clonotypes were presented from each treatment group in **Figure 5C**. Each bubble/circle is indicative of one unique TCR clonotype, and the size of the circle represents how many copies (clones) make up that particular clonotype. Therefore, a larger size circle indicates clonal expansion of that particular clonotype, whereas smaller circles represent clonotypes that failed to expand. SBRT and SBRT/IL-12mRNA treatments resulted in more of the larger sized circles suggesting that the top 1% of CD4 clonotypes underwent increased clonal expansion (**Figure 5C** top; quantified in **D**). In contrast, Untreated and IL-12mRNA had a greater number of smaller circles (poorly expanded TCR clones). The increase in clonal expansion in the top 1% of the CD8 repertoire was more pronounced in the SBRT group with lesser expansion in the SBRT/IL-12mRNA group (yet higher than Untreated and IL12mRNA) (**Figure 5C** bottom; quantified in **D**). Similar results were obtained when the entire downsampled population was examined (**Supplemental Figure 11**). Collectively, these data suggest that SBRT, rather than IL-12, is the driving factor that reshapes the TCR repertoire, which ultimately results in greater clonal expansion of the most dominant clones.

**Figure 5.**
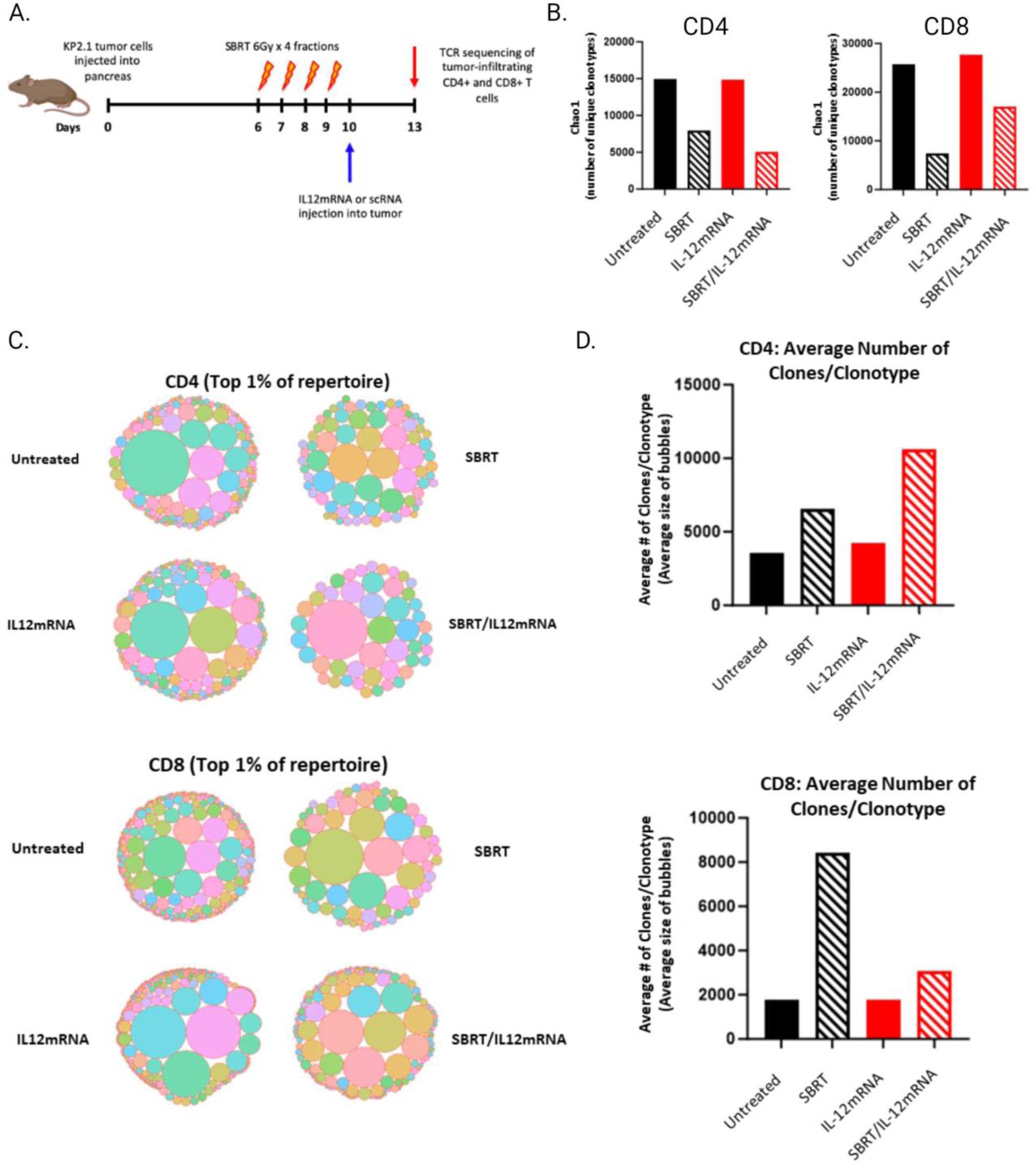
SBRT reshapes the TCR repertoire in tumors. (**A**) Illustration of experimental protocol for TCR sequencing analysis following SBRT/IL-12mRNA treatment: KP2 tumors are injected on Day 0 and SBRT/IL-12mRNA treatment is given as described previously. 3 days after IL-12mRNA/scRNA injection, tumors are harvested, CD4 and CD8 T cells sorted, and their receptors sequenced. (**B**) Chao1 diversity index plot indicating the number of unique clonotypes for each treatment for both CD4 and CD8 T cells. (**C**) Bubble plots for the top clonotypes for each treatment group for both CD4 and CD8 T cells. (**D**) Bar charts depicting the average number of clonotypes for each treatment for both CD4 and CD8 T cells. See materials and methods for further details.

### 2.9 IFNγ mediates SBRT/IL-12mRNA efficacy

Many biological effects of IL-12 are mediated by IFNγ [23]. Additionally, IFNγ, along with IL-12, can skew CD4 T cells towards a Th1 type response that promotes antitumor immunity. This response is typically generated in the dLN, therefore we quantified the IFNγ concentration in the pancreatic dLN and observed a delayed induction of this key cytokine in both the IL-12mRNA alone and SBRT/IL-12mRNA groups (**Supplemental Figure 12A**). Concordantly, a significant expression of the Th1-promoting transcription factor T-bet was observed in the dLN CD4 T cells in the SBRT/IL-12mRNA group (**Supplemental Figure 12B & C**). These data indicate that local combination therapy can polarize immune cells towards an antitumor response in the distal dLN.

We shifted our focus to the tumor and quantified intratumoral IFNγ protein by Luminex in the four treatment groups at multiple time points following therapy. SBRT/IL-12mRNA resulted in elevated concentrations 24 hours after therapy that were sustained for at least 96 hours post therapy (**Figure 6A**). We assessed the production of IFNγ by intracellular flow cytometry of CD45+ cells (**Supplemental Figure 13A**), and IL-12mRNA or SBRT/IL-12mRNA therapy augmented the density of these cells at both time points (**Supplemental Figure 13B**). An immune subset analysis revealed that T cells, and to a lesser extent, NK and myeloid cells, were the predominant intratumoral populations generating IFNγ early, however T cells dominated IFNγ production at later time points (**Supplemental Figure 13C & D**). We focused our analysis on IFNγ+ T cells and showed that CD4 (∼30%) and CD8 (∼20%) had sustained IFNγ production over the 24– and 72-hour timepoints in both the IL-12mRNA and SBRT/IL-12mRNA groups (**Figure 6B**).

**Figure 6.**
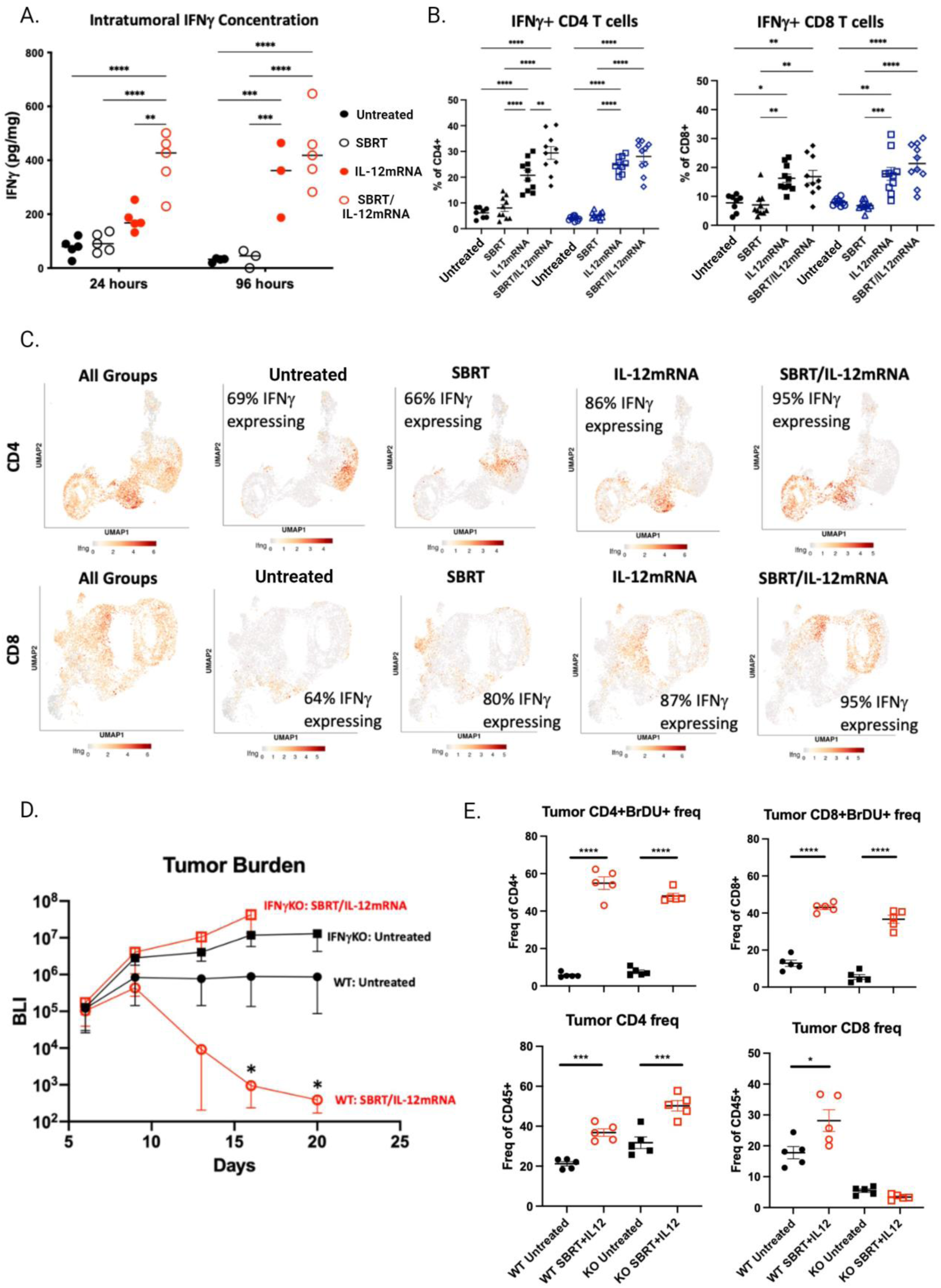
IFNγ expression is induced by SBRT/IL-12mRNA and is essential to treatment efficacy. (**A**) Tumor IFNγ concentration 24 hours and 96 hours post IL-12mRNA injection. **(B)** IFNγ+ CD4 and CD8 T cells at 24 hours (black shapes) and 72 hours (blue shapes) post IL-12mRNA injection. (**C**) UMAPs for CD4 and CD8 T cells for each treatment group depicting levels of IFNγ gene expression. (**D**) SBRT/IL-12mRNA efficacy in IFNγ KO versus wild type mice. (**E**) Proliferation of CD4 and CD8 T cells in IFNγ KO versus wild type mice following SBRT/IL-12mRNA treatment. n=5/group from 2 independent experiments in A. n=10/group from 2 independent experiments. n=5/group in D & E. One-way ANOVA followed by Tukey’s test. Significance against untreated and between treatments are reported. *P<0.05; **P<0.01; ****P<0.0001.

We utilized scRNA-seq data to quantify the frequency and intensity (denoted by heatmap signatures) of each CD4 and CD8 T cell subset (*e.g.* Naïve, Exhausted, etc.) that expressed an IFNγ transcriptomic signature. **Figure 6C (farthest left)** illustrates that the Intermediate population of both CD4 and CD8 T cells has the highest density of IFNγ expression when examining all treatment groups together. When examining each treatment group separately, a lesser IFNγ signature can be observed in the Untreated and SBRT groups (66%), which increases after SBRT/IL-12mRNA treatment (95%) for both CD4 and CD8 T cells. This strong signature could also be appreciated when comparing relative expression levels across treatments, which was considerably higher in both IL-12mRNA and SBRT/IL-12mRNA groups (**Supplementary** Figure 13 **E & F**). Furthermore, the vast majority of all CD4 and CD8 T cells in the SBRT/IL-12mRNA group are expressing IFNγ with the Intermediate group possessing the higher expression when compared to the Activated/Cycling population.

IFNγ was shown to be indispensable in this model as treatment efficacy was completely lost when KP2 tumors grown in IFNγ^-/-^ mice failed to respond to SBRT/IL-12mRNA therapy (**Figure 6D**). We utilized these transgenic mice to assess whether the robust proliferation signature described in Figure 4 was linked to IFNγ production. Either WT or IFNγ^-/-^ mice bearing KP2 and treated as in Figure 1A were administered BrdU four hours before sacrifice and incorporation was examined by flow cytometry. Interestingly, CD4 and CD8 intratumoral T cells from *both* WT and IFNγ^-/-^ mice incorporated BrdU to a similar extent suggesting that intratumoral T cells were able to proliferate without IFNγ (**Figure 6E, upper left and right** respectively). This proliferation still leads to an increased density of intratumoral CD4 T cells regardless of the presence or absence of IFNγ following SBRT/IL-12mRNA therapy (**Figure 6E, bottom left**), however failed to augment the already basally low CD8 T cell density in IFNγ^-/-^ mice (**Figure 6E, bottom right**). Collectively, these data demonstrate that IFNγ, produced predominantly by intratumoral CD4 and CD8 Intermediate and Activated/Cycling T cell populations after SBRT/IL-12mRNA, are required to mediate treatment efficacy.

## 3. Discussion

Chronic antigen exposure along with TME suppressive cues can render T cells to adopt a dysfunctional state known as exhaustion [24]. Immune checkpoint blockade (ICB), which has revolutionized cancer immunotherapy, predominately acts to reverse T cell exhaustion and reinvigorate the adaptive immune response [25]. One striking finding from our work was that SBRT/IL-12mRNA resembled ICB in that intratumoral T cells exhibited an activated/effector phenotype that coincided with an absence of exhausted cells. Although the combination of SBRT and IL-12mRNA resulted in the most profound ratio of activated to exhausted T cells, it was clear that this effect was largely attributed to the presence of intratumoral IL-12 (**Figure 3A & B**). It is well established that IL-12 has direct effects including induction of proliferation [26, 27] and effector function [28], and paramount to this discussion, reversal of T cell exhaustion [29]. Work by Tucker *et al* demonstrated that pre-conditioning adoptively transferred T cells with IL-12 reduced the exhaustion transcription factor Tox and surface expression of Lag3 [30]. Additionally, CAR T cells, which are frequently rendered ineffective due to exhaustion, have been shown to overcome this suppressive phenotype by genetic engineering to constitutively express IL-12 [31]. These previous reports have demonstrated the ability of IL-12 to directly act upon a non-terminally differentiated exhausted T cell and convert it to an effector cell. It is likely that this mechanism is contributing to the T cell conversion in our model. However, another scenario exists in that radiosensitive intratumoral T cells could be reduced by SBRT [32]. These basal state T cells would be an exhausted phenotype and essentially deleted from the TME following radiotherapy, thereby clearing a niche and opening up space for new T cell repopulation. In essence, SBRT may wipe the slate clean allowing for the influx of new T cells that are not yet exhausted and can be strongly polarized towards an activated/effector phenotype by **1)** an inflammatory milieu of cytokines and danger-associated molecular patterns (DAMPs) released by SBRT, and **2)** concentrated IL-12 in the TME. In such a case, the scheduling and order of therapy would be essential requiring SBRT to be given before administration of local IL-12, which would allow for the depletion of exhausted cells, rather than the IL-12-activated T cells.

A robust proliferative signature was observed in the Activated/Cycling T cells from SBRT/IL-12mRNA-treated tumors, which represented over half of all intratumoral T cells. Notably, over 1000 differentially expressed genes were identified in this cluster. A major subset of these genes centered around proliferation and cell division, and were the driving factor in determining UMAP position of the subclustered T cells when analyzed by vector analysis. We were surprised by the limited expression of effector genes in this specific population. One reason for this finding may be linked to the high metabolic demand required for cells undergoing proliferation [33]. Cell division imposes a profound strain on energy consumption as each passage through cell cycle requires the complete doubling of all biomass (macromolecules, lipids, nucleic acids, proteins, etc.) resulting in two daughter cells [34, 35]. This process requires a major reorganization of the cellular metabolic activities allowing the cell to shunt energy into pathways that ultimately support cell division. It is possible that energy is transiently “borrowed” from effector pathways resulting in the observed diminished expression of these genes. Regardless, the SBRT/IL-12mRNA tumors were not devoid of T cells that expressed an effector phenotype. On the contrary, the Intermediate population highly expressed effector associated genes such as IFNγ, granzymes, perforin, etc., but demonstrated a low proliferative signature. The trajectory analysis linked the Activated/Cycling and Intermediate populations together, but intriguingly did not show a terminal population within the activated/cycling cluster. We propose that an external stimuli consisting of IL-12 and other yet unknown factors initiate robust T cell proliferation (the Activated/Cycling population) that when completed, gives rise to a large population of T cells with the capacity to reactivate effector function to elicit antitumor activity (the Intermediate population). This concept is intriguing as it suggests that a unique milieu of SBRT/IL-12mRNA-induced factors can support *intratumoral* T cell expansion; an anatomical location not typically considered as being hospitable to support T cell division.

Historically, IL-12 and IFNγ have been linked as potent inducers of T cell proliferation. Some publications attribute this characteristic as a direct effect of IL-12 through a STAT4-mediated mechanism [26, 36], whereas others highlight the importance of IFNγ stimulating T cell division [37, 38]. In our model, the frequency of intratumoral proliferating T cells (as assessed by BrdU incorporation) was essentially identical between WT and IFNγKO mice. These data suggest that IL-12 alone (or IL-12 induced factors) were driving T cell proliferation independent of IFNγ. Importantly, all treatment efficacy was lost in IFNγKO mice even though proliferation was intact. This suggests that T cell proliferation without IFNγ renders these T cells completely ineffective in this model. Therefore, what essential role is IFNγ having? The answer likely lies in the difference between the Activated/Cycling and the Intermediate population as described above. The Activated/Cycling cells are dividing but express little IFNγ and a muted effector phenotype, whereas IFNγ is more abundant, along with effector genes, in the Intermediate population. The IFNγKO mice have proliferating T cells, but poor effector function, emphasizing the importance of this Intermediate population in mediating tumor cell killing. Other antitumor roles for IFNγ presumably exist. The vast majority of IFNγ was produced by T cells allowing for autocrine stimulation (or paracrine from CD4 T cells) of cytolytic capabilities of CD8 T cells. IFNγ is well established to act on intratumoral cells such as tumor, stromal, endothelial and myeloid cells often promoting antitumor activity [39]. Distally in the draining lymph node, we demonstrated Tbet induction; evidence of a strong induction to a Th1 response driven by IL-12-induced IFNγ. Collectively, T cell proliferation is important to expand intratumoral T cell populations, however IFNγ licenses them to become effector T cells and ultimately eliminate the tumor.

Although SBRT has been shown to downstage locally-advanced or borderline resectable tumors in PDAC patients [40], it rarely results in a complete clinical response or long-term survival. We observed similar results in our preclinical model as SBRT alone had minimal effects on tumor burden and survival, however, when combined with IL-12, could completely eliminate disease. Therefore, what is the contribution of SBRT to this therapy? We have demonstrated both preclinically [22, 41] and clinically [7] that SBRT induces immunogenic cell death of tumor cells with subsequent release of DAMPs such as HMGB1, HSP70, and calreticulin. In cancer, DAMPs bind to cellular cognate receptors to sense damage/danger in the TME, which, in turn, stimulates proinflammatory cytokine release such as Type I IFNs [8]. This cascade of events is essential to initiate antitumor immune responses. Dendritic cells are the predominate immune subset that react to DAMPs, however “bystander” T cells also respond to these molecules through TLR, and this promotes T cell activation in a TCR-independent manner [42, 43]. There is evidence in our model that SBRT is also stimulating a response to DAMPs as a strong Type I IFN signature is observed in macrophages, monocytes, and DCs (**Supplemental Figure 8).** In addition to providing the essential ingredients required to initiate an antitumor immune response, SBRT is also reshaping the TCR repertoire. The optimal antitumor T cell response would be to generate clonal expansion against numerous tumor antigens. We have shown that SBRT increased expansion of the top 1% clonotypes, which likely translates into extended coverage of the TCR repertoire against more tumor antigens. We propose that SBRT exposes and releases novel tumor antigens that would otherwise not be recognized by the immune system without this treatment modality. Typically, this is the point where the expanded repertoire of T cells would possess TCRs that specifically recognize tumor cells, only to be suppressed by the PDAC TME via checkpoint molecules and/or induction of exhaustion. However, intratumoral IL-12 prevents this suppression and instead, promotes robust proliferation and effector function of these tumor-specific T cells directly at the tumor site. In essence, the combination of SBRT and IL-12 evokes T cell proliferation against a diverse set of tumor antigens, which do not become exhausted and go on to eliminate the pancreatic tumor.

The data presented here demonstrate that local administration of SBRT and IL-12 essentially propels every cell in the TME towards an antitumor state. This “sledgehammer” approach is capable of rejecting primary tumors, but also stimulates a systemic response that eliminates distal hepatic metastases. SBRT is already an approved standard of care for PDAC patients. Intratumoral administration of antitumor therapeutics is an emerging field as investigators explore non-systemic means to introduce drugs directly to the tumor. Endoscopic ultrasound guided biopsy is routinely used in PDAC [44], and can be repurposed to deliver immunotherapeutics such as IL-12mRNA. Collectively, this work represents evidence that even the recalcitrant PDAC TME can be overpowered when the correct combination of therapies (in this case SBRT and IL-12mRNA) are administered locally.

## 4. Materials and Methods

### 4.1. In vivo animal studies

C57BL/6J and B6.129S7-Ifng^tm1Ts^/J (IFNg^-/-^) mice were purchased from The Jackson Laboratory. P48-Cre^+/-^; Tp53^L/L^ and LSL-Kras^G12D^ mice were obtained from Dr. Aram Hezel and crossed to generate the LSL-Kras^G12D^; Tp53^L/L^; P48-Cre; (KPC) genotype [13, 14]. All experiments were approved by the University Committee on Animal Resources and were performed in compliance with the NIH and University of Rochester approved guidelines for the care and use of animals.

### 4.2. Cell Lines

KP2 (derived from KPC mice) and KCKO (derived from KC mice; LSL-Kras^G12D^; P48-Cre) cell lines stably express firefly luciferase and were provided by Dr. David DeNardo [15]. Both cell lines were cultured in MAT/P (US patent No. 4.816.401) supplemented with 5% fetal bovine serum (Hyclone) and 1% penicillin/streptomycin (GIBCO) and incubated at 37°C, 5% CO_2_.

### 4.3. Orthotopic tumor mouse model

Mice were anesthetized using vaporized isoflurane (Scivena Scientific) and a 10-mm laparotomy incision was made to expose the spleen and pancreas. KP2.1-luc (2.5×10^4^) or KCKO-luc (5×10^4^) cells were resuspended in a 1:1 PBS:Matrigel (Corning) solution and injected into the tail of the pancreas (40ul volume). Two 4-mm titanium fiducial clip markers (Horizon) were implanted adjacent to the Matrigel bubble to assist in SBRT targeting. The peritoneal cavity was then sutured with 4-0 VICRYL SH-1 (eSutures) and skin was stapled using 9-mm wound clips (Reflex).

### 4.4. Metastasis model

Mice were anesthetized using vaporized isoflurane and a 20-mm laparotomy incision was made to expose the abdominal cavity. The hepatic portal vein was exposed and 1×10^5^ KP2.1-luc cells in 100ul of PBS were injected into the vein using a 30-gauge Hamilton syringe to establish hepatic metastasis. Concurrently, primary pancreatic tumors were injected as described above. The peritoneum was sutured and skin stapled as before.

### 4.5. Radiation Therapy

All radiation was delivered using the Small Animal Radiation Research Platform (SARRP, XStrahl) equipped with a CT scanning device (Muriplan software) as described previously [16]. Briefly, tumor-bearing mice were anesthetized with vaporized isoflurane and transferred to the SARRP. Pancreatic tumors were identified by a CT image based on two titanium fiducial clips placed on either side of the tumor at the time of tumor injection. SBRT was administered to the tumor following a dosing schedule of 4 fractions each consisting of 6 Gy radiation on days 6-9 post-implantation.

### 4.6. IL-12mRNA injection

IL-12mRNA and scRNA lipid nanoparticles were provided by Moderna Inc. and their synthesis and design have been described previously [12]. Briefly, the IL-12mRNA encoded sequences for wild-type mouse IL12α and IL12β subunits joined by a polypeptide linker to generate a stable IL12p70 fusion protein as reported elsewhere [18]. ScRNA (control for IL-12mRNA) was modified to exclude an initiating codon and therefore is non-translating. Twenty-four hours after the final SBRT fraction (day 10 post tumor implantation), mice were anesthetized with vaporized isoflurane and a 10-mm laparotomy incision was made to expose the pancreas and tumor. ScRNA or IL-12mRNA (0.5ug in 25ul) were injected intratumorally using a 30-gauge Hamilton syringe.

### 4.7. Bioluminescent Imaging

*In vivo* tumor growth was measured using the *In Vivo* Imaging Systems (IVIS, Perkin Elmer). Tumor-bearing mice were anesthetized and injected subcutaneously (s.c.) with D-luciferin (2.5mg/100ul PBS, Invitrogen). Mice were placed in a right lateral recumbent position and a series of 12 consecutive images were taken at 2-minute intervals. BLI (p/sec/cm^2^/sr) was calculated within matching (circular) regions of interest (ROI) manually placed over tumors. Peak intensity was recorded for each tumor and used as a readout of tumor burden.

### 4.8. Rechallenge model

Mice were injected with orthotopic tumors and treated with SBRT and IL-12mRNA as described above. Six weeks after IL-12mRNA injection (4 weeks after loss of BLI signal), mice were rechallenged systemically with an injection of 1×10^5^ KP2-luc cells through the hepatic portal vein as described above. Additionally, treatment-naïve aged-matched mice were also injected at the same time as controls.

### 4.9. Flow Cytometry

Tumors were mechanically dissociated on day 1, 3 and 7 post IL-12mRNA/scRNA injection (day 11, 13 and 17 post inoculation) and digested in 30% collagenase (Sigma) for 30 minutes at 37°C. Homogenates were then passed through a 70 µm filter to generate a single cell suspension which was redispersed in PBS containing 1% bovine serum albumin and 0.1% sodium azide. This suspension was then stained for various markers.

To assess immune populations in tumors, single cell suspensions were stained with the following: Ghost Dye 510 (Tonbo), CD4-BUV395 (RM4-5, BD), CD11b-AF700 (M1/70, Biolegend), F4/80-APC (BM8, Life Technologies), Ly6C-PEcy7 (HK1.4, Biolegend), CD8-PEcy5 (53-6.7, BD), NK1.1-PECF595 (PK136, BD), CD45-PerCPcy5.5 (30-f11, BD), CD19-BV786 (ID3, BD), CD11c-BV711 (HL3, BD), Ly6G-BV650 (IA8, Biolegend), TCRγδ-BV605 (GL3, Biolegend), IFNγ-PE (XMG1.2, Biolegend), and Foxp3-APC (FJK-16s, eBioscience). For intercellular staining of IFNγ BD Cytofix/Cytoperm Plus kit (BD) was used.

For T cell proliferation analysis, 1mg of 5-bromo-2-deoxyuridine (BrdU) was injected intraperitoneally four hours prior to tumor harvest on day 13 post tumor inoculation (3 days post IL-12mRNA/scRNA treatment). Cells were then stained according to BrdU APC staining kit instructions (BD).

All samples were run on a LSRII Fortessa (BD Bioscience) and analyzed using FlowJo software.

### 4.10. Single-Cell RNA-Sequencing

For scRNA-seq, 2 KP2.1 tumors were pooled per treatment group at 3 days post IL-12mRNA/scRNA injection (day 13 post tumor inoculation), minced and digested in collagenase (Sigma) for 30 minutes at 37°C. Single cell suspensions were stained with Ghost Dye 510 (Tonbo). Live singlet cells were sorted using the FACSAria II (BD) using a 100um nozzle and provided to the University of Rochester Genomics Research Center for RNA purification, sequencing and analysis. ScRNA-seq libraries were generated using Chromium Single-Cell (10x Genomics) according to manufacturer’s instructions with a target cell capture of 10k cells per sample and 100k reads per cell.

Libraries were sequenced on Illumina NovaSeq 6000 sequencer. Raw sequencing data were processed using Cell Ranger (v6.0.1) and aligned to mouse reference genome GRCm38.

Quality control was performed using Seurat (version 4.2.0) by removing cells with low quality RNA based on high mitochondrial gene content (Percent mt > 20%), or low unique gene counts (nFeature < 200). To avoid doublets, cells with greater than 6500 feature counts were also excluded. The remaining cells were normalized, anchor-point analysis performed, and merged according to Seurat pipeline guidelines to form a combined seurat object for downstream analysis.

Cell clustering and differential analysis was performed via the Seurat pipeline using the Louvain algorithm. Clusters were annotated using known marker genes and manual inspection of cluster-specific gene expression patterns based on positive expression and minimum percentage expression by cluster greater than 25%. Secondary automated classification was performed using sc-Type for cross comparison. Differential expression analysis was conducted using a Wilcoxon rank-sum test using an adjusted p-value (Bonferroni false discovery rate) threshold of 0.05. Cluster comparisons were made as specified in the text either between manual cluster classification for all cells in that cluster of the merged seurat object and or by cluster classification subset by condition of origin. For subset analysis, CD4 and CD8 seurat objects were subset on CD3e > 1.0 AND CD4 > 0.5 OR CD8a > 1.0 respectively. Pseudotime trajectories were generated using Monocle3 (Version 1.3.1) using manually identified naïve T cells as the root population for each seurat object.

### 4.11. TCR-Sequencing

For TCR sequencing (TCR-seq), 5 KP2.1 tumors were pooled per treatment group 3 days post IL-12mRNA/scRNA injection (day 13 post tumor inoculation), minced and digested in collagenase (Sigma) for 30 minutes at 37°C. Single cell suspensions were stained with Ghost Dye 510 (Tonbo), CD45-PerCPcy5.5 (30-f11, BD), CD4-BV605 (RM4-5, BD) and CD8-PEcy5 (53-6.7, BD). Cells were sorted into 2 populations, CD4+ and CD8+, using a FACSAria II (BD) using a 100um nozzle. Cells were immediately lysed in RLT Plus buffer (Qiagen) and delivered to the University of Rochester Genomics Research Center for sequencing and analysis.

TCR-seq data was run through MiXCR (MiLabs) using a pipeline for analysis of enriched targeted TCR libraries. The pipeline performed alignment of raw sequencing reads, assembly of aligned sequences into clonotypes, and output of the resulting clonotypes into tab-delimited files. The final list of clones was further analyzed using Seurat and Immunarch in R. TCR clones were downsized randomly to the lowest number of clones in the four groups (Untreated, SBRT, IL-12mRNA, SBRT/IL-12mRNA). Clonotypes were defined as the set of all TCR clones with the same CDR3 nucleic acid sequence. High clonality indicates fewer overall clonotypes, with each clonotype having a higher frequency. Low clonality indicates the even distribution of clonotypes across the repertoire. Chao1 diversity was calculated using Immunarch in R. Bubble plots displaying the top 1% of clonotypes in each TCR repertoire were generated using Seurat. The size of each bubble corresponds to the number of clones with that clonotype. The average size of the bubbles in each plot were calculated and plotted.

### 4.12. Antibody Depletion

Depleting antibodies were administered (200ug in 100ul PBS) subcutaneously beginning on day 5 post KP2.1 tumor implantation (one day before SBRT begins) and continued every three days for a total of 7 doses. The antibodies (Bio X Cell) administered were rat isotype control (clone 2A3), rat anti-mouse CD8a (clone 53.6.7), rat anti-mouse CD4 (clone GK1.5), anti-mouse NK1.1 (clone PK136), mouse isotype control (clone C1.18.4).

### 4.13. Immunohistochemistry

3 days post IL-12mRNA/scRNA injection (day 13 post-implantation) KP2.1 tumors were harvested, paraffin embedded, and 5μm tumor sections were stained with hematoxylin and eosin (H&E). Slides were analyzed in a blinded fashion by a board-certified pathologist. Images are representative of 3 individual tumors for each treatment group.

### 4.14. Luminex assay

KP2.1 tumor, draining lymph node (dLN), non-draining lymph node (ndLN), and liver were harvested 1 day or 4 days post IL-12mRNA/scRNA injection (day 11 or 14 post inoculation), homogenized and digested in Cell Lysis Buffer #2 (R&D Systems) containing protease inhibitors on ice for 1 hour. The samples were centrifuged at 14,000 rpm for 20 min at 4°C and supernatants were collected. Magnetic Luminex Assays were preformed using a Mouse Premixed Cytokine/Chemokine Multi-Analyte kit (R&D Systems) according to manufacturer’s instructions. Microplates were run on a Bio-Plex 200 system (Biorad). Total protein concentrations for each sample were calculated using Pierce BCA Protein Assays (ThermoFisher Scientific) and used for analyte normalization into pg/mg protein values.

### 4.15. ELISA and qPCR

*In vitro* cultured cells, KP2, mouse embryonic fibroblasts (MEFs), naïve CD4/CD8 lymphocytes and peritoneal lavage macrophages, were treated with IL12mRNA or scRNA for 24 hours. Supernatants were assayed for IL-12 protein with the Murine IL-12 Standard ELISA Development Kit (Peprotech) according to manufacturer’s instructions. RNA was isolated from the cells using the RNeasy Plus Mini Kit (Qiagen), cDNA was then generated with the iScript cDNA Synthesis Kit (Bio-Rad) according to manufacturer’s instructions. The protocol from IDT, Prime Time qPCR Assay reactions (Premixed primer and probes) was followed to assay for a unique sequence in IL-12mRNA using the CFX96 Real-Time System (C1000 Thermocycler, BioRad).

### 4.16. Statistical analysis

GraphPad Prism 9 software was used for all statistical analyses and P values of <0.05 were considered significant. BLI growth curves used the geometric mean of maximum photon emissions for each timepoint and treatment group from a single ROI. Growth curve significance were assessed using ANOVA followed by Dunn’s multiple comparison test. For survival analysis, Mantel-Cox tests were performed. Rechallenge studies were analyzed by Mann-Whitney test. Reported significance was against the untreated group, unless otherwise stated.

### 4.17. Data availability

The data generated in this study are available upon request from the corresponding author. Transcriptomic data in this study is publicly available in Gene Expression Omnibus (GEO) at GSE242111.

