## Supplementary data for "Local Delivery of SBRT and IL12 by mRNA Technology Overcomes Immunosuppressive Barriers to Eliminate Pancreatic Cancer"

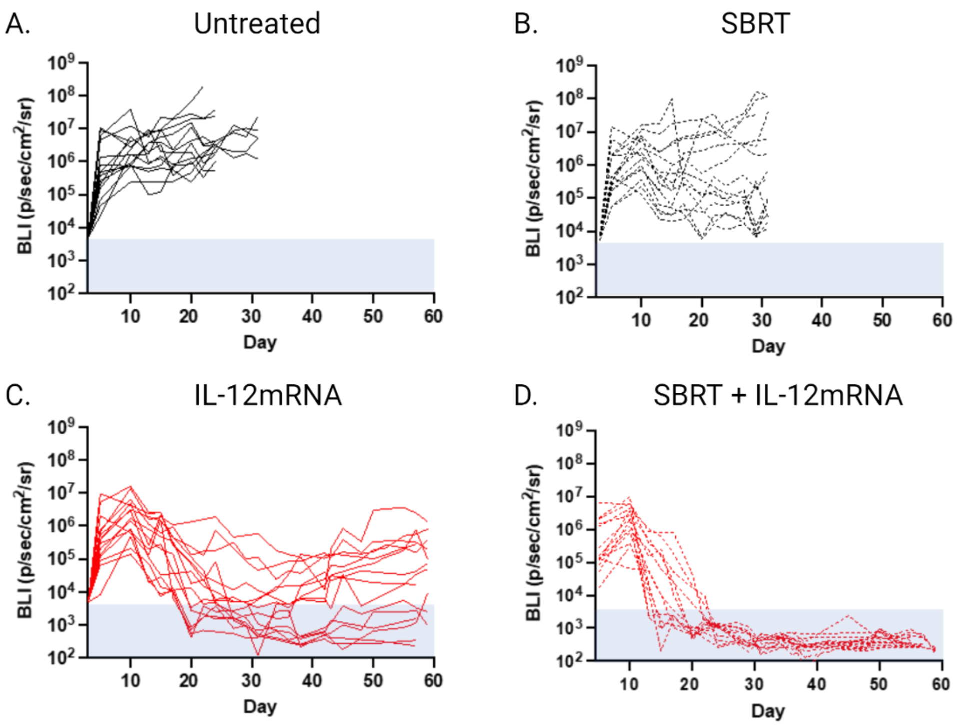


**Supplementary Figure 1. Individual BLI growth curves for primary tumors treated with SBRT and IL-12mRNA along with single treatment controls.** Tumor burden over time was recorded by weekly BLI measurements and individual growth curves plotted for unirradiated, scRNA treated (**A**), SBRT and scRNA treated (**B**), unirradiated and IL-12mRNA treated (**C**) and SBRT and IL-12mRNA treated (**D**) mice. Grey area denotes baseline BLI levels indicative of cures. Data represents n=15/group from 3 independent experiments.

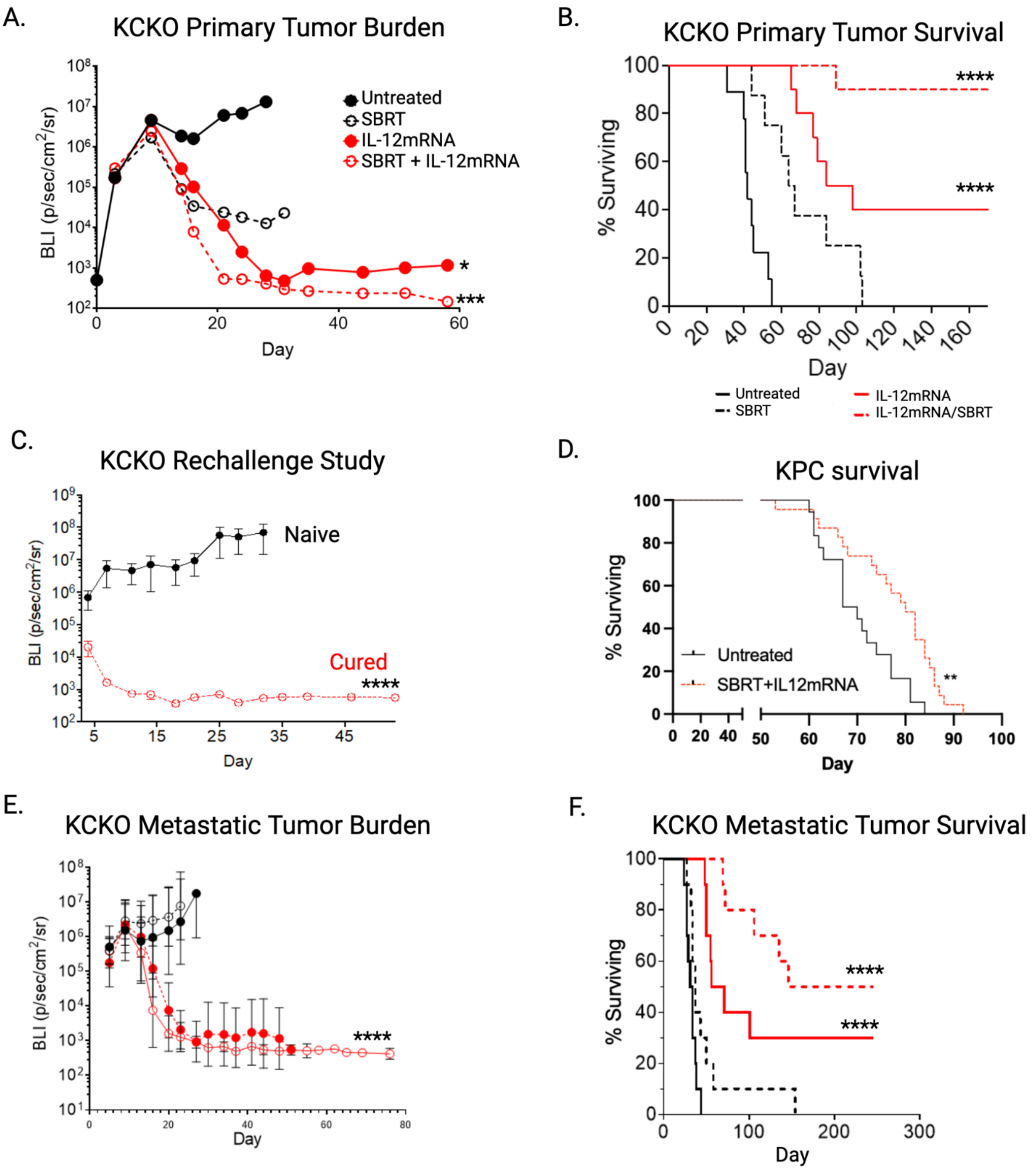


**Supplementary Figure 2. Combination treatment controls tumor burden and improves survival in multiple PDAC mouse models.** (**A**) KCKO cells were injected orthotopically and treated with or without SBRT (6Gy/day over 4 consecutive days) followed by an intratumoral injection of IL-12mRNA or scRNA 1 day after the final fraction of radiation. Tumor burden was assessed by BLI measurements. (**B**) Survival of KCKO model followed for >160 days. (**C**) Mice cured of KCKO tumors following SBRT and IL-12mRNA treatment for 150 days were rechallenged with a hemi-spleen injection of KCKO-Luc to induce hepatic metastases. Liver tumor burden was assessed by IVIS. Aged-matched naïve mice were used as controls. (**D**) KPC mice were assessed for tumor at 5 weeks of age and fiducial clips inserted accordingly. SBRT (6G/day over 4 consecutive days) followed by an intratumoral injection of IL-12mRNA or control scRNA 1 day after the final fraction of radiation was administered and survival monitored. (**E**) KCKO primary tumors were implanted along with liver metastasis and primary tumors were treated with SBRT and IL-12mRNA. Tumor burden was measured by BLI. (**F**) Survival of metastatic KCKO model. n=10/group from 3 independent experiments in A & B; n=5 naïve and n=7 cured in C; n=18 for Untreated and n=23 for SBRT + IL-12mRNA in D. n=10 in E and F. *p<0.05;**p<0.01; ***p,0.001 ****p<0.0001.

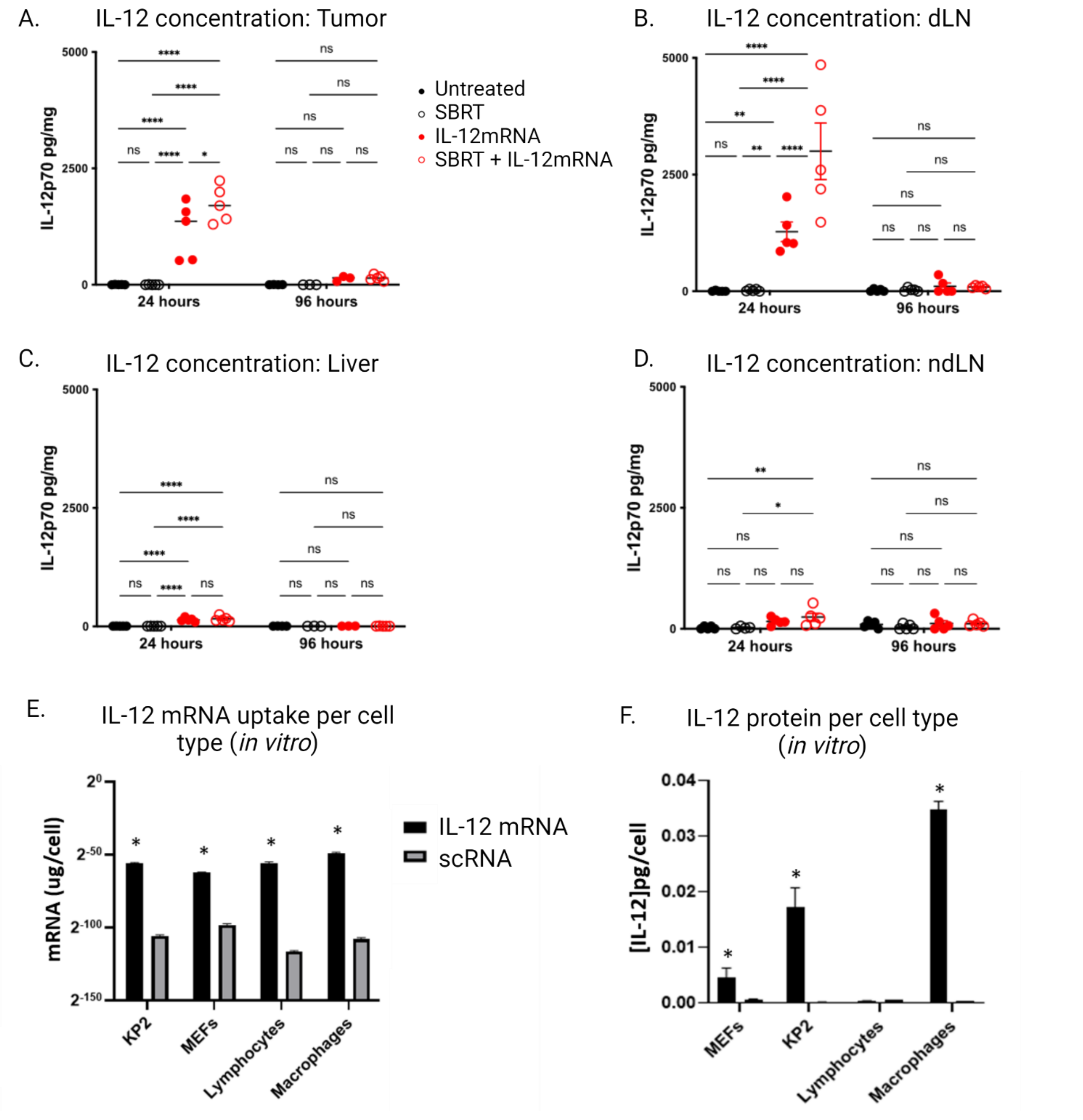


**Supplemental Figure 3. Assessment of IL-12 concentration following treatment.** Mice were inoculated with KP2 tumor cells and treated as detailed in **Figure 1A**. 24 and 96 hours after intratumoral injection of IL-12mRNA or scRNA, tissues were removed, homogenized, and assayed for IL-12p70 protein by Luminex. IL-12 concentration was determined following standardization to total protein (pg/mg) in (**A**) primary pancreatic tumors, (**B**) pancreatic tumor-draining lymph node, (**C**) liver, and (**D**) non-draining (inguinal) lymph node. Y-axis is kept uniform to compare across tissues. (**E**) Cultured cells consisting of KP2, mouse embryonic fibroblasts (MEFs), naïve CD4/CD8 lymphocytes, and peritoneal lavage macrophages were treated with IL-12mRNA or scRNA *in vitro* for 24 hours and assayed for a unique sequence in IL-12mRNA by qPCR (**E**) or IL-12 protein by ELISA (**F**). Values are standardized to number of total cells recovered at endpoint. n=5 mice/group in A-D; In E&F, each sample well was performed in triplicate and experiment repeated three times. Analyzed by one-way ANOVA followed by Tukey's test in A-D; student t-test in E&F comparing IL-12mRNA to scRNA in each group. $^{*}p<0.05$; $^{**}p<0.01$; $^{****}p<0.0001$.

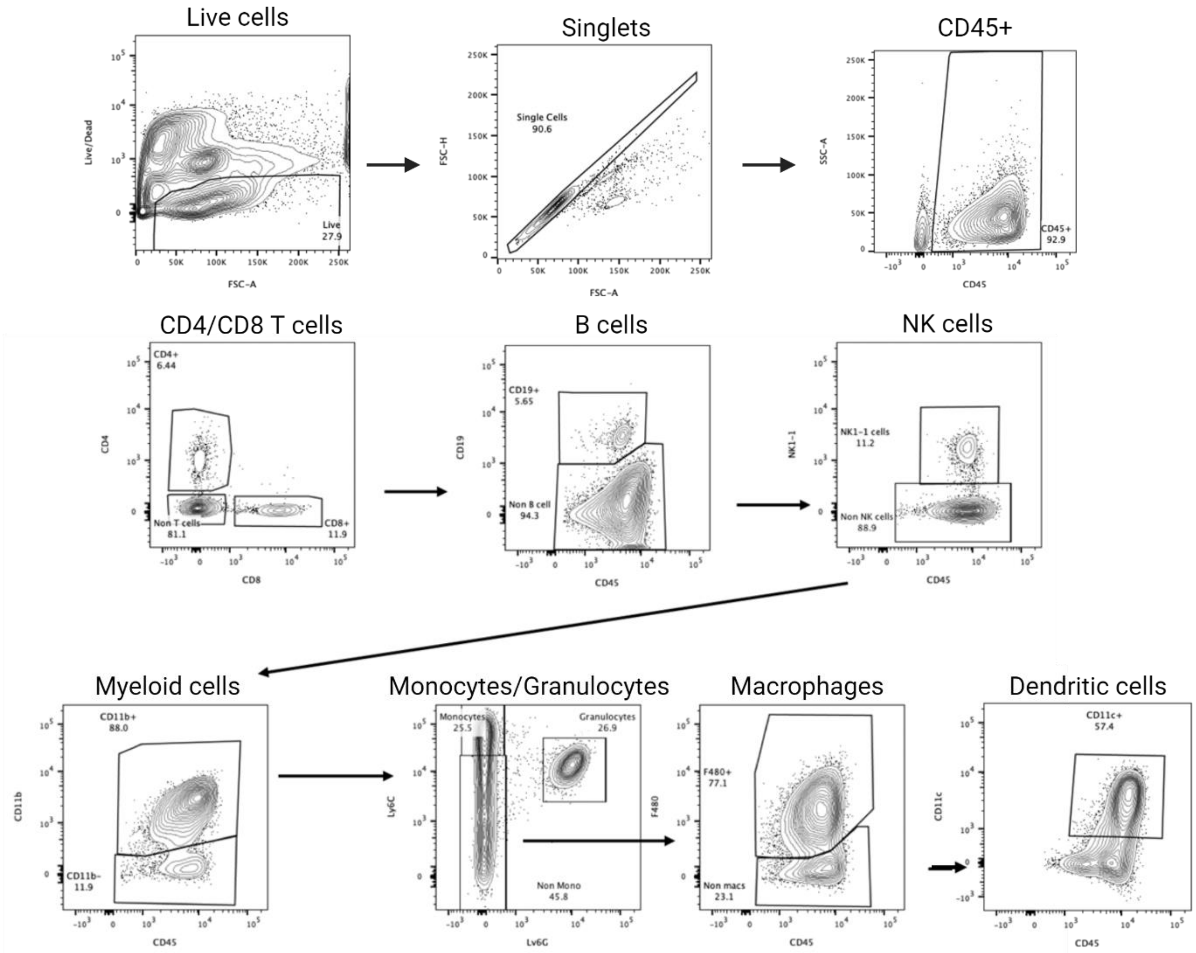


**Supplementary Figure 4: Gating strategy for determining immune subsets.** Tumors were treated as described in **Figure 1A** and were harvested at 24 and 72 hours post IL-12 mRNA/scRNA injection and stained for various markers to determine changes to immune subsets. Live cells (Ghost dye-) were gated against FSC-A and doublets were removed. CD45+ cells were gated and used to determine CD4+ and CD8+ T cells. Cells negative for CD4 and CD8 were used to gate B cells (CD19+), NK cells (NK1.1+) and myeloid cells (CD11b+). Myeloid cells were used to identify monocytes (Ly6C+), granulocytes (Ly6G+Ly6C+), macrophages (Ly6C-F480+) and dendritic cells (F480-CD11c+).

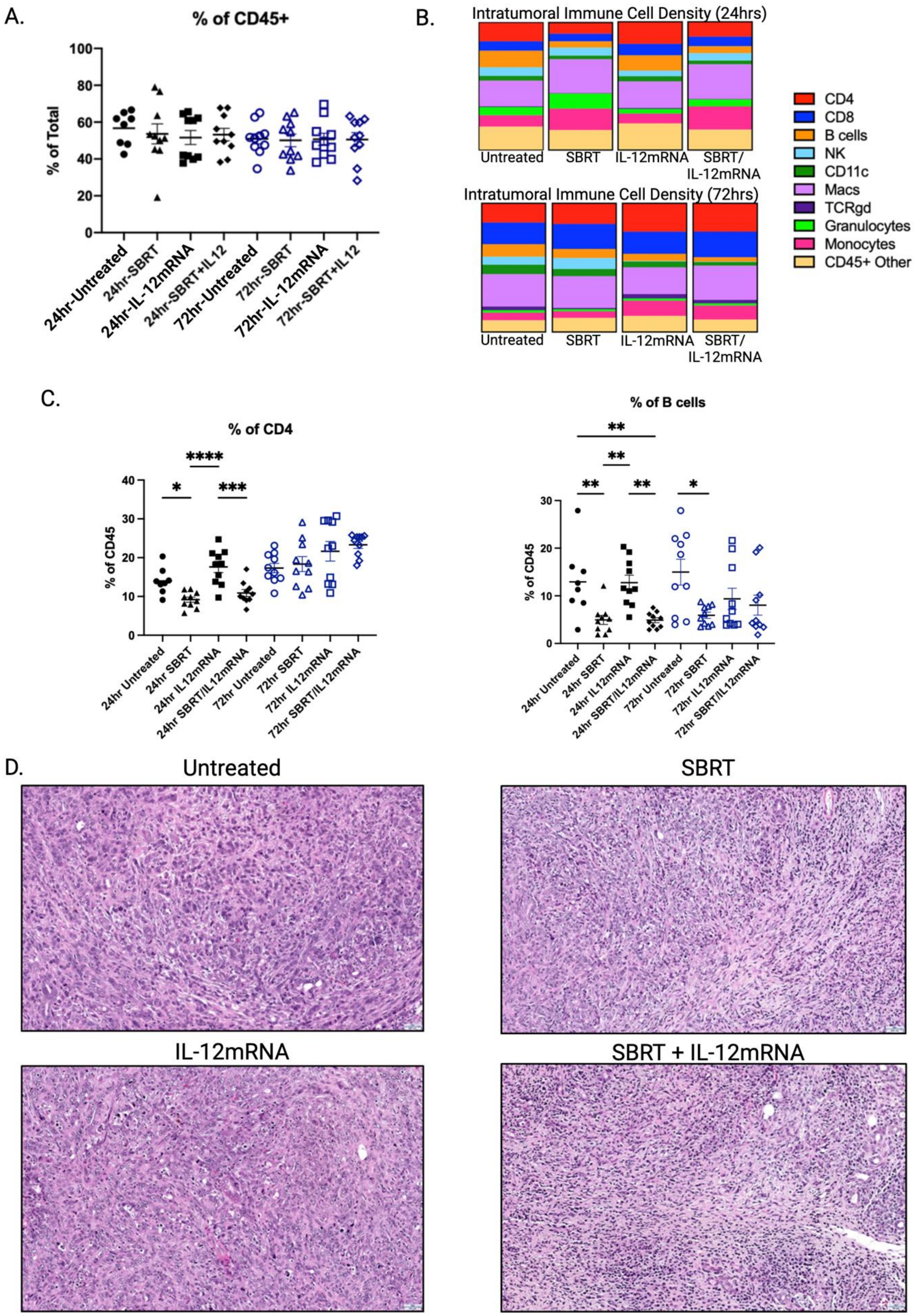


**Supplementary Figure 5. Effects of SBRT/IL-12mRNA on the tumor microenvironment.** (**A**) No changes in levels of CD45+ cells were observed across all treatments at 24 and 72hrs post IL-12mRNA/scRNA injection. (B) Flow cytometry analysis of immune subsets across all treatments and time points. (C) Frequency of CD4 T cells and B cells 24hrs and 72hrs post IL-12mRNA/scRNA injection for each treatment group (**D**) Representative H&E images of tumors 72hrs post IL-12mRNA/scRNA injection (20X magnification). n=8-10 for A, B and C. Analyzed by one-way ANOVA followed by Tukey's test in C. *p<0.05; **p<0.01; ****p<0.0001.

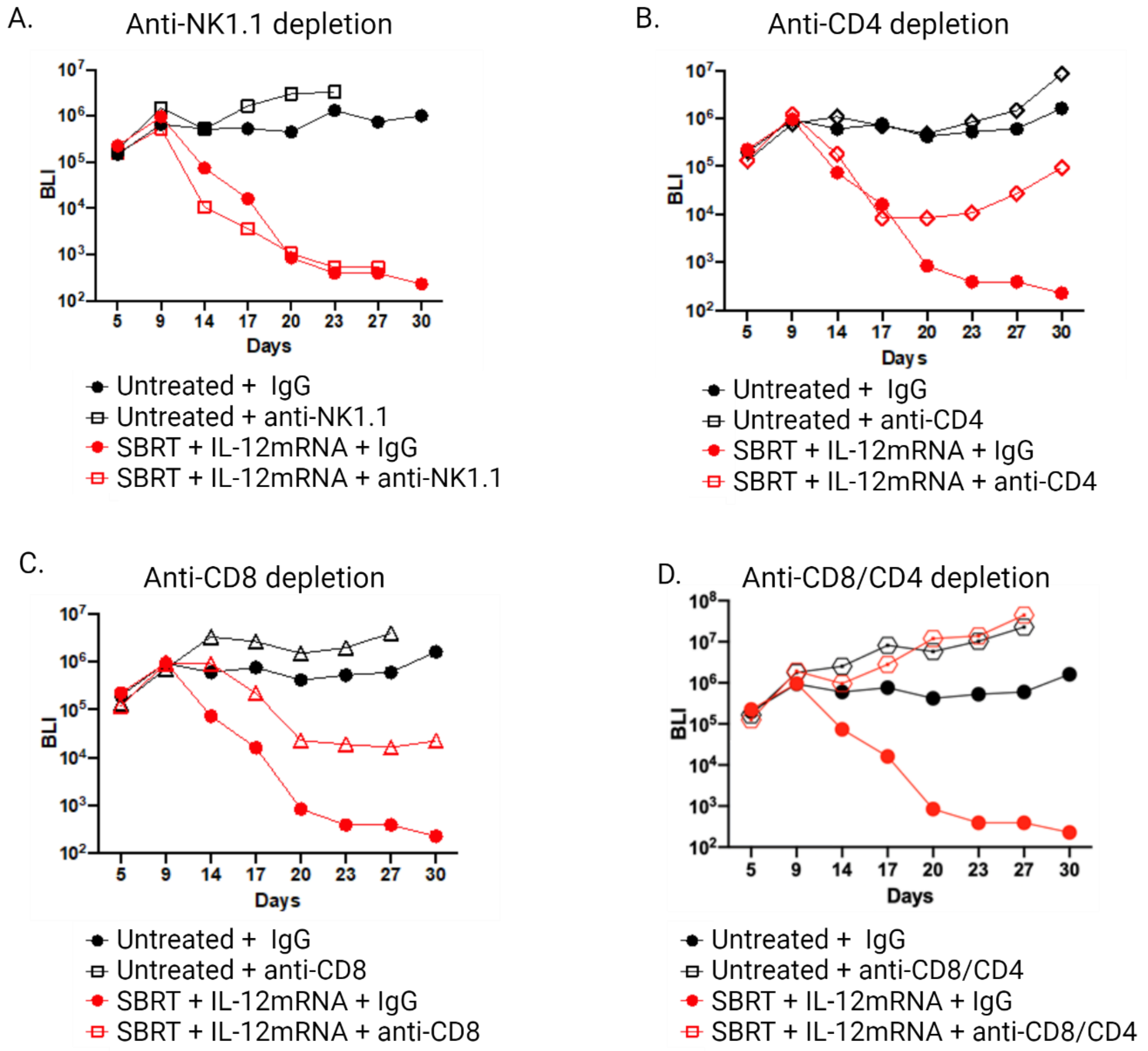


**Supplementary Figure 6. BLI growth curves following antibody depletion of effector cell populations during and after SBRT/IL-12mRNA treatment**. (**A**) NK cell depletion with isotype control for untreated and SBRT/IL-12mRNA treated mice. (**B**) CD4 T cell depletion with isotype control for untreated and SBRT/IL-12mRNA treated mice. (**C**) CD8 T cell depletion with isotype control for untreated and SBRT/IL-12mRNA treated mice. (**D**) CD4 and CD8 T cell depletion with isotype control for untreated and SBRT/IL-12mRNA treated mice. In all cases, depletions began at Day 5 post tumor implantation (1 day prior to the first fraction of SBRT) and continued every 3 Days for a total of 7 doses. n=10 for each group.

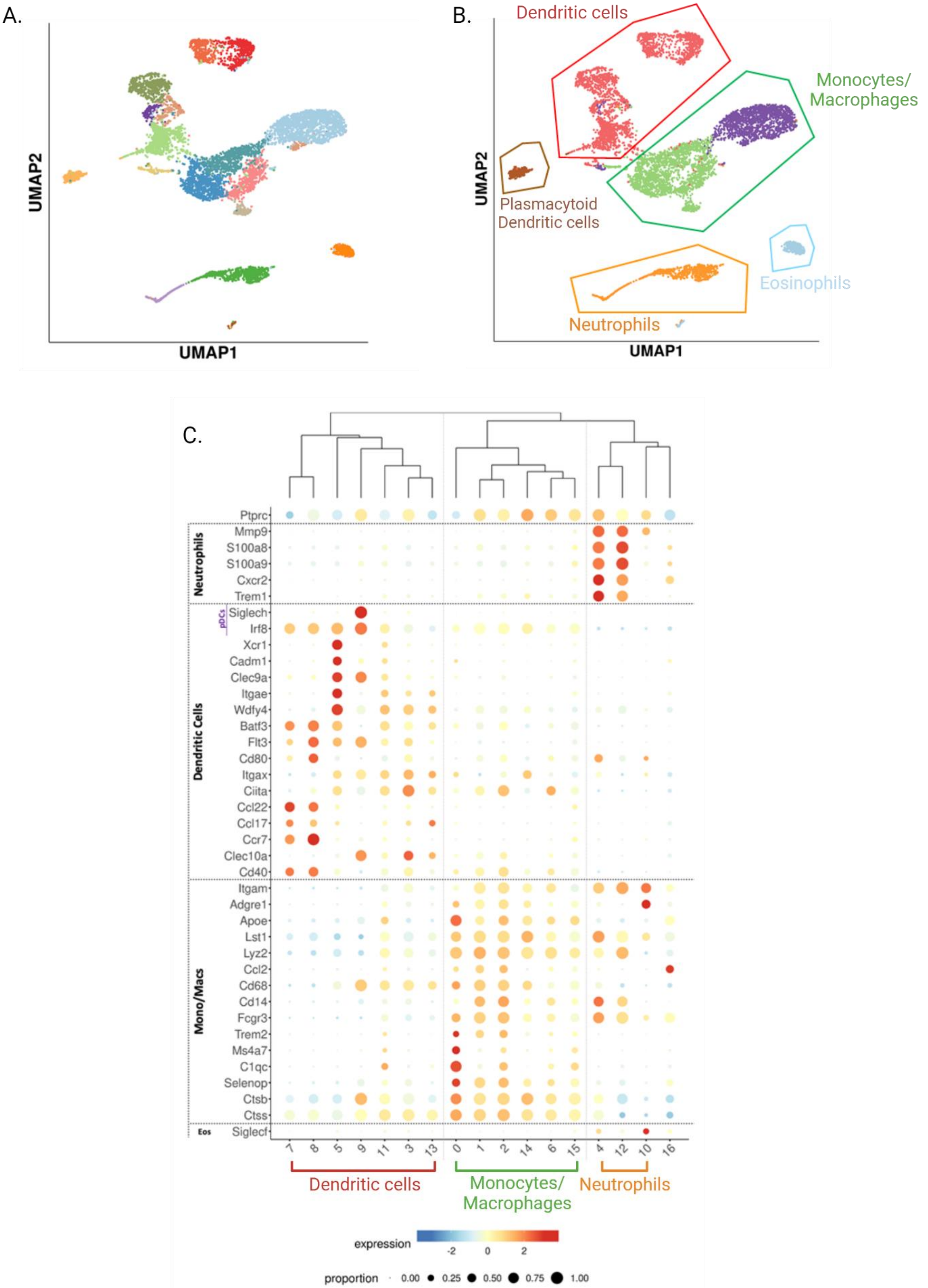


**Supplementary Figure 7. Identifying myeloid populations with scRNA-seq.** (**A**) Combined UMAP for all myeloid cells for each treatment. (**B**) Identified populations of dendritic cells, monocytes/macrophages, neutrophils and eosinophils based on canonical markers. (**C**) Dot plots of canonical markers highlighting the different populations of myeloid cells.

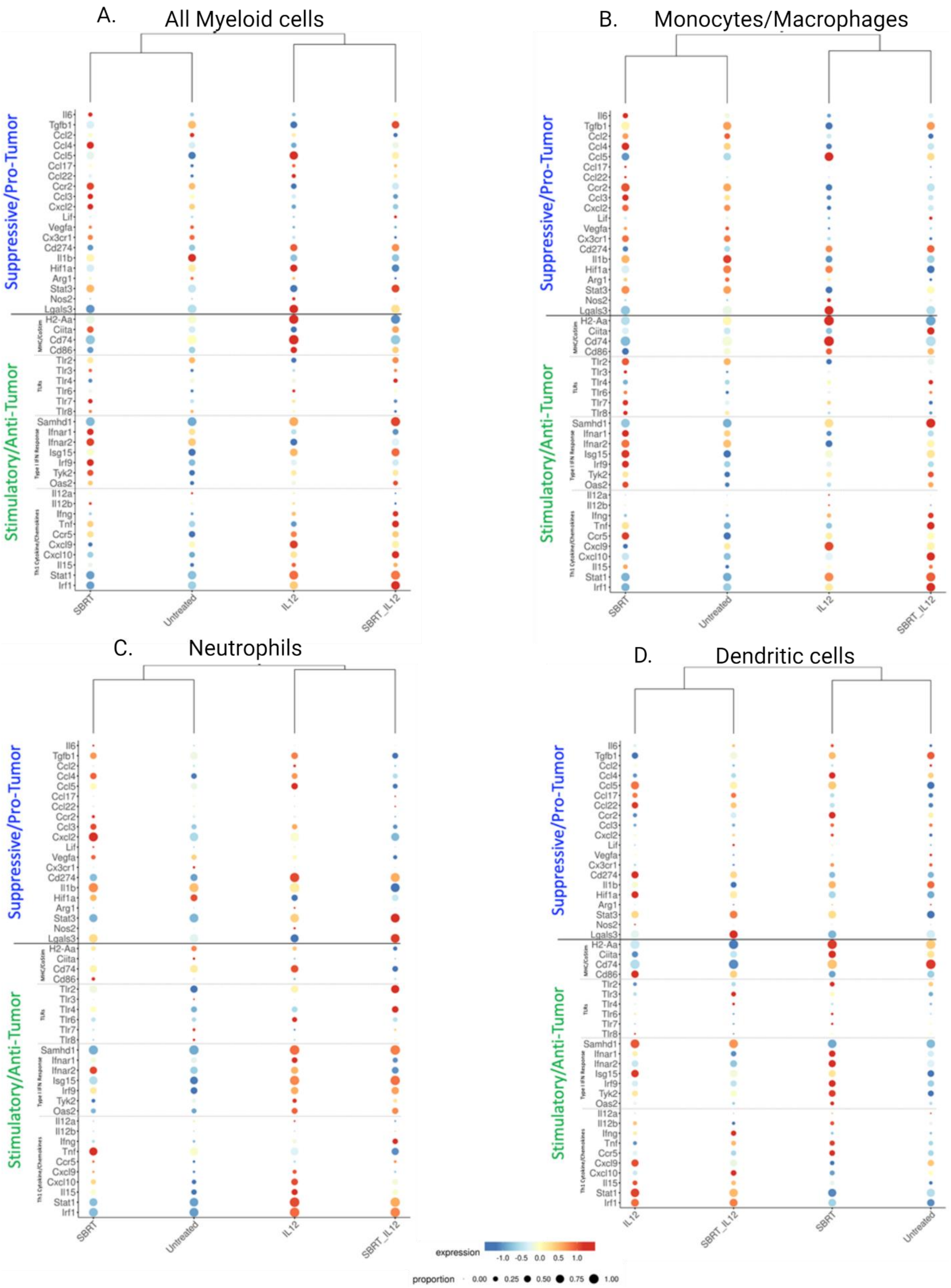


**Supplementary Figure 8. Expression of immunostimulatory and immunosuppressive genes in myeloid populations following treatment.** Canonical immunostimulatory and immunosuppressive gene expression in (**A**) All Myeloid cells, (**B**) Monocytes and Macrophages, (**C**) Neutrophils and (**D**) Dendritic cells following treatment.

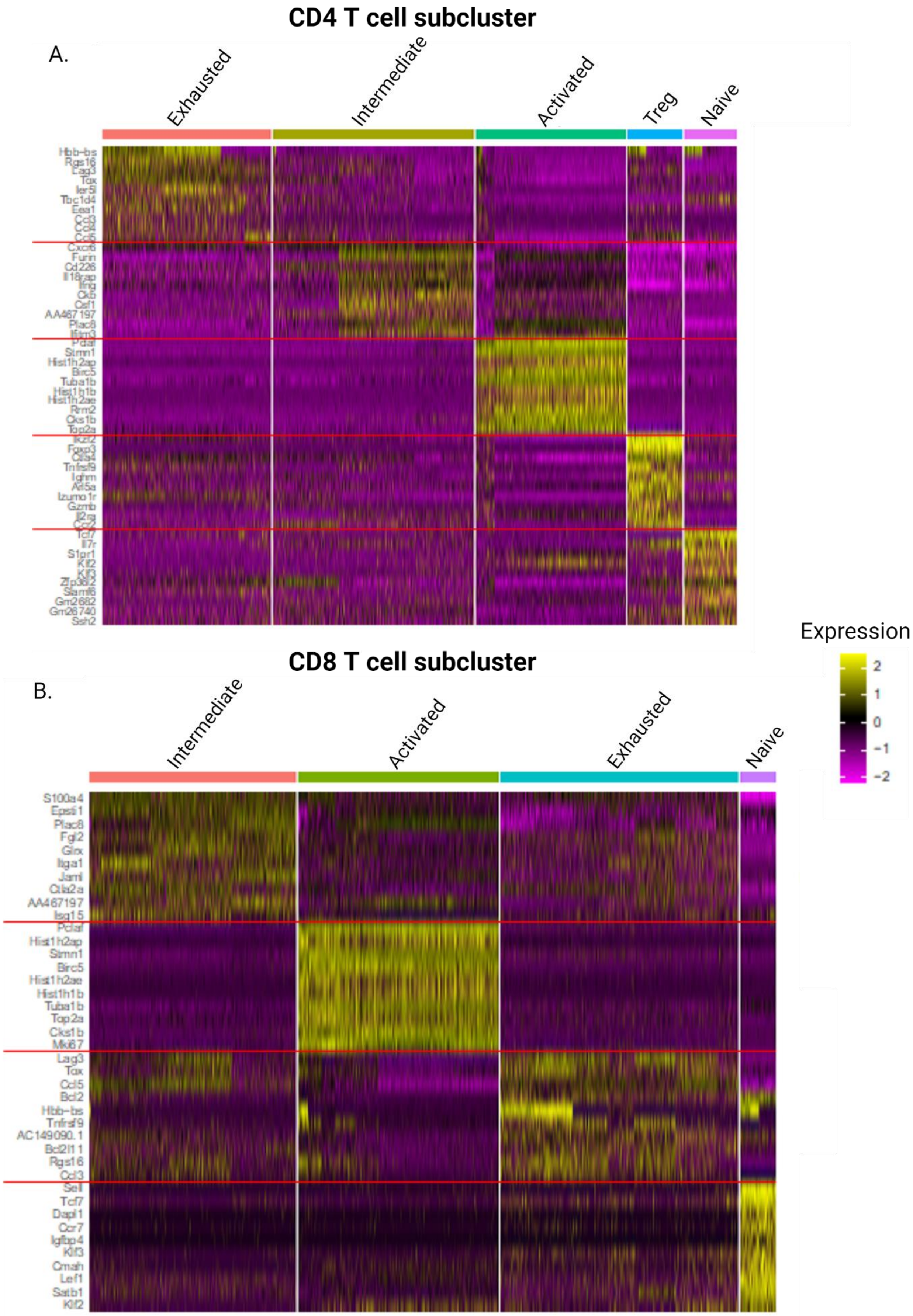


**Supplementary Figure 9. Differentially expressed genes for each subcluster of CD4 and CD8 T cells.** Top 10 differentially expressed genes for each subcluster of CD4 (**A**) and CD8 (**B**) T cells are presented in heat maps.

A.

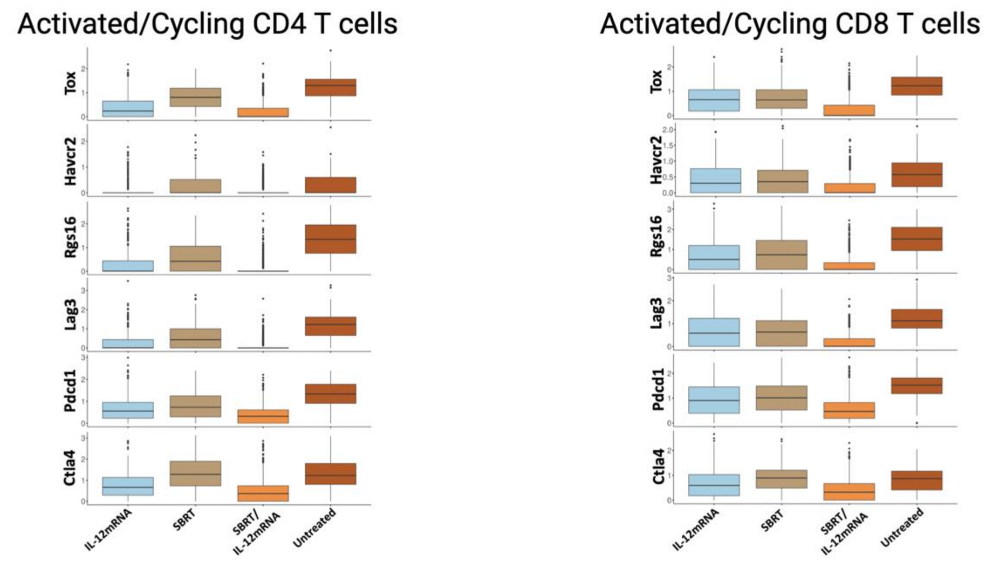


B.

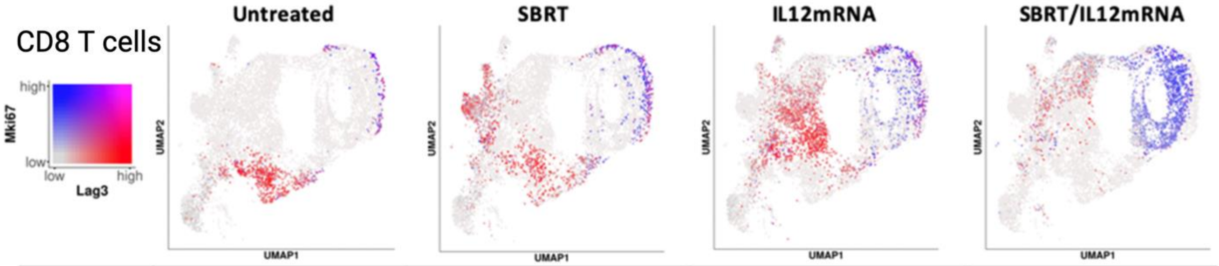


| Expression | # of Cells | % | # of Cells | % | # of Cells | % | # of Cells | % |
|---|---|---|---|---|---|---|---|---|
| Dual | 300 | 24 | 567 | 29 | 714 | 27 | 782 | 30 |
| Lag3 | 561 | 44 | 800 | 41 | 1309 | 50 | 444 | 17 |
| Mki67 | 18 | 1 | 172 | 9 | 211 | 8 | 902 | 35 |
| None | 387 | 31 | 401 | 21 | 413 | 14 | 437 | 17 |

C.

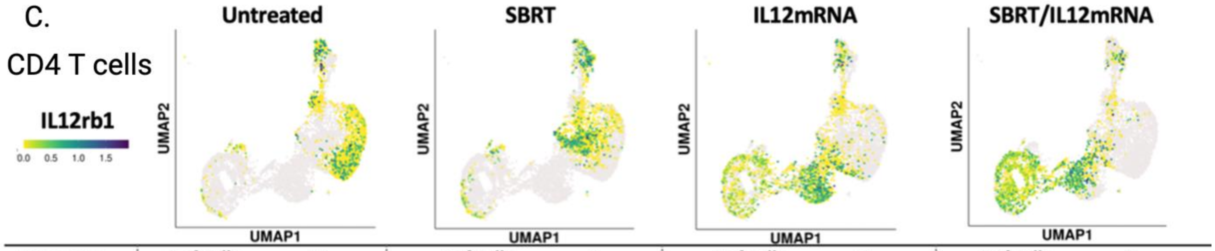


| Expression | # of Cells | % | # of Cells | % | # of Cells | % | # of Cells | % |
|---|---|---|---|---|---|---|---|---|
| IL12rb1 | 313 | 15 | 426 | 25 | 710 | 34 | 856 | 40 |
| None | 2121 | 85 | 1727 | 75 | 2089 | 66 | 2120 | 60 |

D.

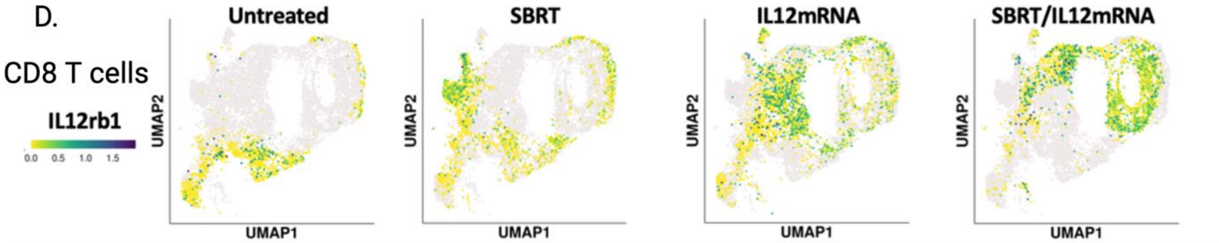


| Expression | # of Cells | % | # of Cells | % | # of Cells | % | # of Cells | % |
|---|---|---|---|---|---|---|---|---|
| IL12rb1 | 156 | 12 | 435 | 22 | 806 | 30 | 947 | 37 |
| None | 1266 | 88 | 1940 | 78 | 2647 | 70 | 2565 | 63 |

**Supplementary Figure 10. Expression of key genes in CD4 and CD8 T cells post treatment.** (**A**) Boxplot depicting exhaustion marker expression in activated/cycling CD4 and CD8 T cells following treatment. (**B**) A marked reduction in exhaustion marker Lag3 is accompanied by an upregulation of proliferative marker Mki67 after SBRT/IL-12mRNA treatment. (**C** & **D**) IL-12mRNA treatment induces an upregulation of its receptor subunit IL-12rb1 in CD4 and CD8 T cells in a potential positive feedback loop.

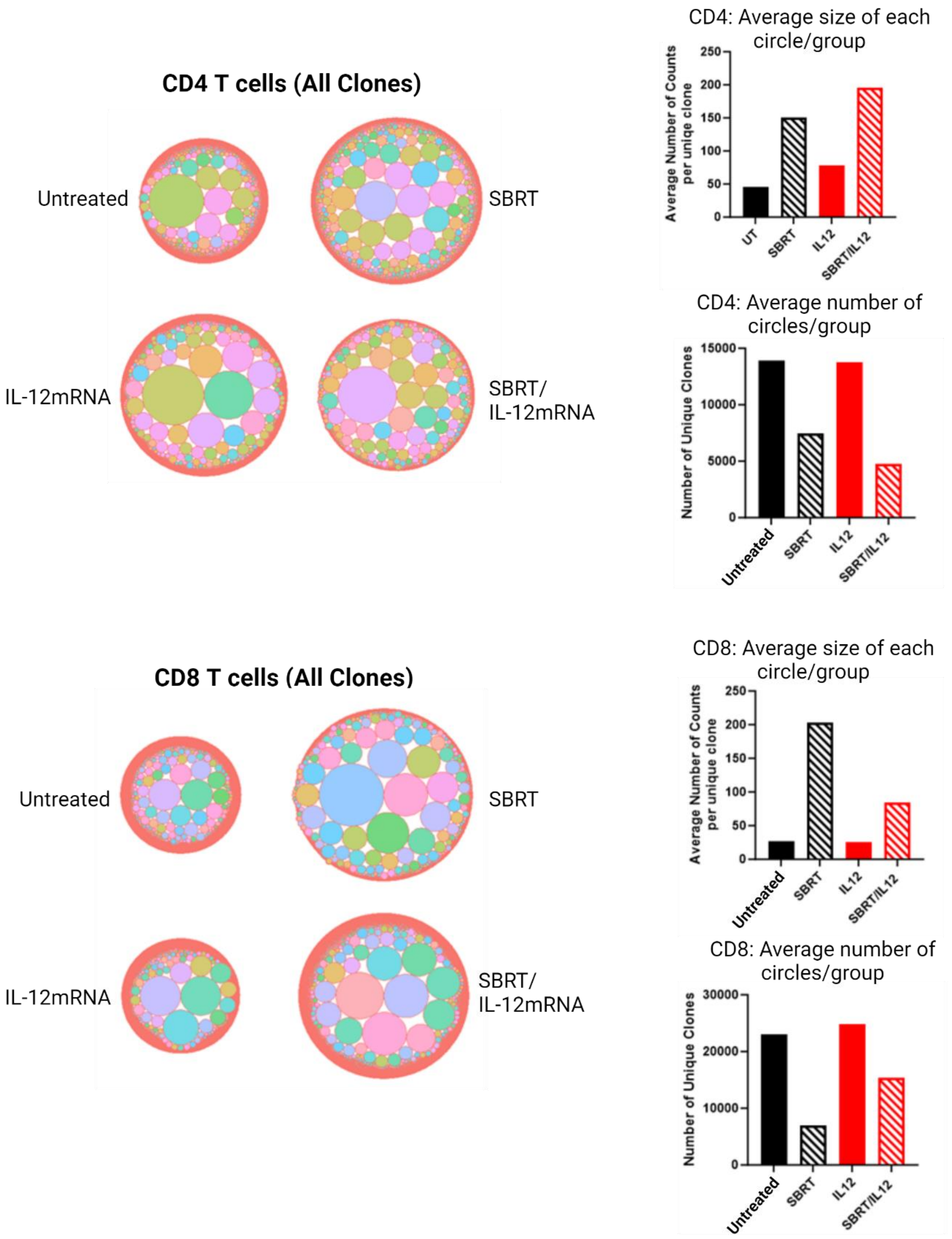


**Supplementary Figure 11. Downsampled population of CD4 and CD8 T cell clonotypes.** Bubbleplots and accompanied bar chats depicting the downsampled size of T cell receptor clones and their relative abundance are detailed for both CD4 and CD8 T cells.

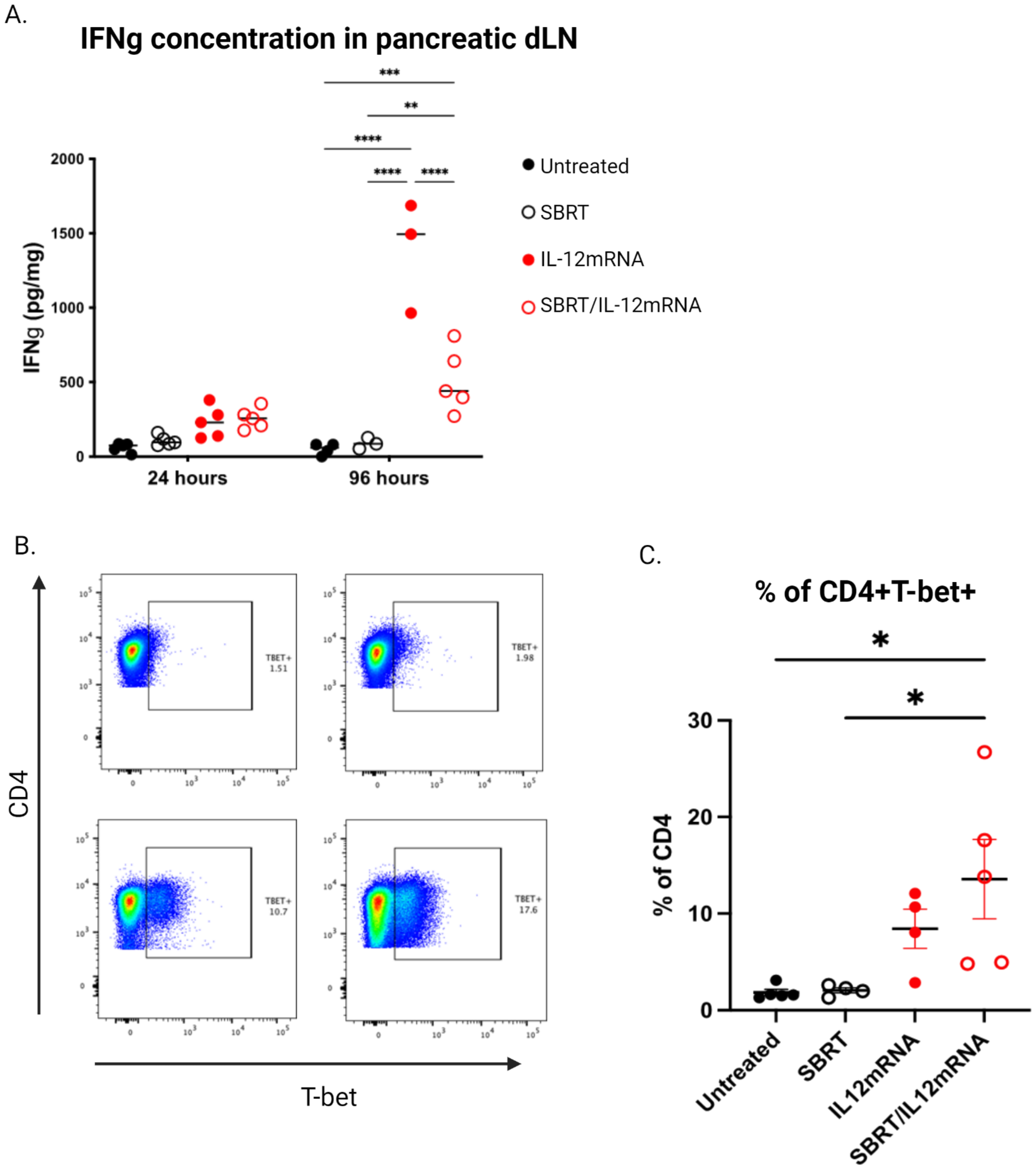


**Supplementary Figure 12. Quantifying IFNγ and T-bet expression in pancreatic draining lymph node following treatment.** (**A**) IFNγ is significantly increased in the pancreatic draining lymph node 96 hours post IL-12mRNA/scRNA injection. (**B**) Representative flow plots for T-bet staining of CD4 T cells. (**C**) Significant increases of T-bet+ CD4 T cells in the draining lymph node were were determined at 72hrs post IL-12mRNA/scRNA injection. Both A and C were analyzed by one-way ANOVA and Tukey's test. Significance is compared against the untreated and within treatments. n=5 in both cases. *p<0.5; **p<0.01; ****p<0.0001.

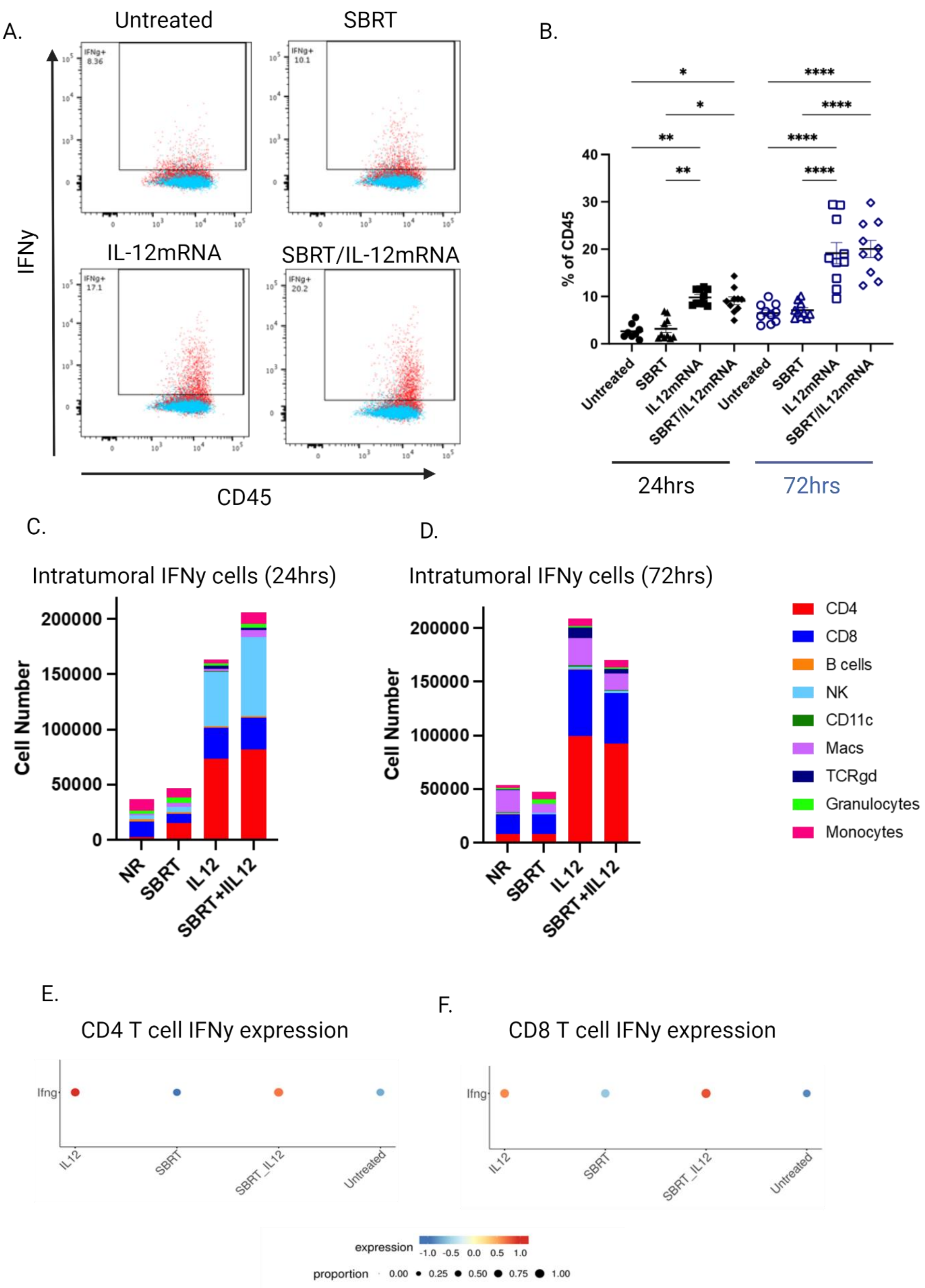


**Supplementary Figure 13. Expression of IFNγ in the tumor microenvironment following treatment.** (**A**) Representative flow plots for IFNγ staining of CD45+ cells from the tumor microenvironment. Blue events indicate FMO control, red events correspond to IFNγ staining. (**B**) Tumor levels of IFNγ+ CD45+ cells 24hrs and 72hrs post IL-12mRNA/scRNA injection. (**C & D**) Relative numbers of IFNγ+ intratumoral immune cells 24hrs and 72hrs post IL-12mRNA/scRNA injection. (**E & F**) Gene expression of IFNγ+ by CD4 and CD8 T cells is mediated by IL-12mRNA. B was analyzed by one-way ANOVA followed by Tukey's test. *p<0.05; **p<0.01; ****p<0.0001.
